# Volatile diacetyl triggers rapid *fmo-2* expression through DHAP-glycerol shunt induction

**DOI:** 10.64898/2026.07.01.735788

**Authors:** Marco Giorda, Anamitra Sen, Martin S Denzel

**Affiliations:** Cambridge Institute of Science, Altos Labs, Cambridge, United Kingdom; University of Cologne, Cologne, Germany; Max Planck Institute for Biology of Ageing, Cologne, Germany

## Abstract

*C. elegans* can locate ephemeral food sources by sensing bacterially-derived volatiles. However, whether these olfactory cues can also trigger anticipatory nutrient-responsive programs remains unknown. Using unbiased transcriptomics and metabolomics, we discover that fasting worms exposed to the volatile food cue diacetyl rapidly induce expression of enzymes in the DHAP-Glycerol shunt, a metabolic pathway at the intersection of glycolysis and glycerolipid biosynthesis. We demonstrate that this pathway, known to be activated by glucotoxicity and hyperosmotic stress, is also modulated by food availability. Shunt induction requires the MDT-15 transcription factor and drives rapid metabolic remodeling, characterised by the accumulation of glycerol and phosphatidylglycerols. When food deprivation persists, the shunt’s activity triggers expression of the dietary restriction marker *fmo-2*, promoting thermotolerance and food-seeking behaviours. Our findings reveal that volatile diacetyl can rapidly modulate gene expression in *C. elegans*, providing new insights into the DHAP-Glycerol shunt function in metabolic homeostasis.

## Introduction

Anticipatory food responses allow animals to prepare for food absorption in response to food sensing. Olfactory and gustatory cues have been shown to stimulate production of digestive enzymes and hormones, most notably insulin^1^. Recent research has extended this concept from physiological secretions to the level of gene expression. Food perception can alter hepatic gene expression in mice as early as thirty minutes after exposure, inducing responses that overlap with those induced by refeeding after food deprivation^2^. Similarly, exposing food-deprived *Drosophila* to thirty minutes of vinegar odor triggers transient expression of genes encoding glucagon-like adipokinetic hormone and insulin-like peptides^3^. This work has focused on specific organs^2^ or individual genes^3^, leaving whole-genome and systemic organismal responses largely unexamined.

*C. elegans* represents an ideal, yet underexplored, model organism for examining anticipatory food responses. It naturally inhabits environments like decaying vegetable matter, where its bacterial food source is found in ephemeral patches^4^. To thrive in this unstable ecological niche, worms have evolved flexible developmental programs^5^ and a highly developed chemosensory system. This allows them to detect food by identifying byproducts of bacterial metabolism^6^. One of these byproducts is diacetyl, a volatile α-diketone produced by lactic acid bacteria (LAB) to dispose of pyruvate excess, typically resulting from citrate metabolism^7^. Diacetyl serves as one of the most well-characterized food signals for *C. elegans*; it is generated by the LAB that worms consume in their natural habitat^8^ and is a powerful chemoattractant across various concentrations^6^. While several studies have addressed the role of *C. elegans* olfaction in regulating long-term processes like healthspan and lifespan^9, 10, 11^, the immediate effects of exposure to food odors have not been widely explored. In particular, it is not known whether food olfactory cues can trigger anticipatory responses preparing worms for feeding-related stress.

Glucose excess can overwhelm glycolysis and mitochondrial respiration, leading to an accumulation of reducing equivalents (NADH) and glycolytic intermediates that further disrupt glycolytic flux and contribute to cellular stress^12^. Glucotoxicity-related redox stress particularly leads to accumulation of intermediates upstream of GAPDH (Glyceraldehyde 3-phosphate dehydrogenase)^13^, including GAP (glyceraldehyde 3-phosphate) and DHAP (dihydroxyacetone phosphate)^14, 15^. To relieve this block, cGPDH (cytosolic glycerol-3-phosphate dehydrogenase) reduces DHAP into Gro3P, regenerating NAD+ in the process. Significantly, cGPDH activity has been identified as a conserved protective mechanism in conditions of hypoxia and ETC dysfunction in yeast, *C. elegans*, mice and human cells^16^. GPDH-1 has also been shown to modulate *C. elegans* lifespan in high-glucose conditions^17^.

However, Gro3P accumulation itself can drive excessive lipid synthesis and result in metabolic dysfunction^18^. To deal with this secondary stress, Gro3P is converted into glycerol, a less reactive solute that can be readily secreted from the cell. The conserved enzyme G3PP (glycerol-3-phosphate phosphatase), also known as PGP (phosphoglycolate phosphatase) in mammals or PGPH (PGP homolog) in *C. elegans,* hydrolyses the phosphate group from Gro3P to produce glycerol^18^. This reaction has been named “glycerol shunt”, and protects both mice^19^ and *C. elegans*^20^ against glucotoxicity.

Together, GPDH and G3PP activities convert DHAP to Gro3P and Gro3P into glycerol, forming the DHAP-Glycerol shunt. The DHAP-Glycerol shunt from the alga *Chlamydomonas reinhardtii*, which operates both reactions from a single enzyme, has been recently applied to *in vitro* and *in vivo* mammalian models to relieve metabolic stress caused by hypoxia, mitochondrial dysfunction and ethanol^21^. In *C. elegans*, components of the DHAP-Glycerol shunt are mostly studied in the context of hyperosmotic stress^20,22^, where glycerol is accumulated as the primary protective osmolyte. Significantly, expression of *gpdh-1* is utilised as an indicator for the hyperosmotic stress response^23^.

We exposed food-deprived *C. elegans* to the olfactory food cue diacetyl to investigate temporal dynamics and crosstalk between acute gene expression and metabolic responses. We established an experimental model for volatile exposure to a 1% solution of diacetyl. This is the diacetyl concentration most attractive to worms^6^ and has been previously adopted for chronic exposure studies^24^. We found that fasting worms exposed to diacetyl rapidly induce expression of *gpdh-1* and other DHAP-Glycerol shunt enzymes, a pathway otherwise repressed during food deprivation. DHAP-Glycerol shunt activation in the absence of food also caused a strong secondary upregulation of FMO-2, a marker of dietary restriction^25^ and key regulator of endogenous metabolism^26^. These findings reveal a novel function of volatile food cues in the rapid modulation of the nutritional status of an animal.

## Results

### Volatile diacetyl exposure induces rapid expression of nutrient-modulated DHAP-Glycerol shunt pathway

To identify the earliest and most direct effects of diacetyl exposure, we opted for a transcriptomics time course at high temporal resolution with time points between 5 and 90 minutes (**Fig. 1a**). These time points are in line with the speed of transcriptional anticipatory food responses in other organisms^2, 3^. Consistent with what has been done in the *Drosophila* and mouse studies, we used food-deprived animals for the transcriptomics analysis, as well-fed animals would presumably be less responsive to a food signal.

**Figure 1.**
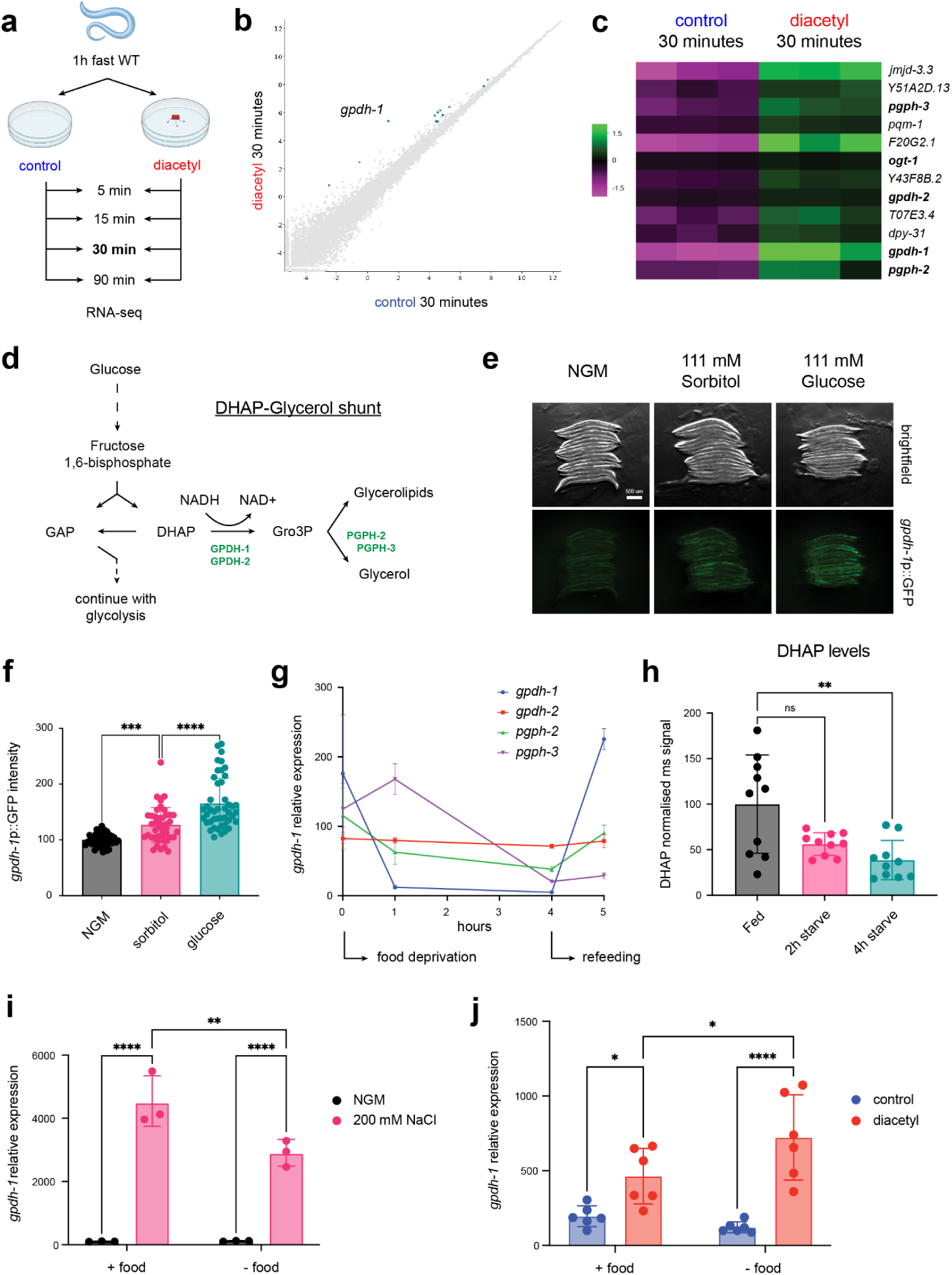
Volatile diacetyl exposure induces rapid expression of nutrient-modulated DHAP-Glycerol shunt pathway. **a**, Experimental setup for measuring gene expression effects of diacetyl exposure in food-deprived worms, comparing control food deprivation (no volatile) to food deprivation in the presence of volatile diacetyl (1% solution in ethanol, pipetted on plate lid). Three biological replicates collected for RNA-seq over a ninety-minute time course. **b,** Scatter plot showing differential gene expression between control and diacetyl groups at 30 minutes of exposure. **c,** Heatmap showing differential gene expression between control and diacetyl groups at 30 minutes of exposure, highlighting genes involved in glycerol production. **d,** Schematic depicting DHAP-glycerol shunt pathway, connecting glycolysis to synthesis of glycerol and glycerolipids. Highlighted in green are enzymes upregulated by diacetyl. **e,** Representative images of *gpdh-1* transcriptional reporter worms raised on NGM, sorbitol or glucose supplemented plates. Images shown are from one experiment out of three biological replicates. Additional replicates are available in Figure 1 - Source data e. **f,** Quantification of reporter induction in experiment from (e). Data is pooled from all three replicates (n = 45 worms per condition). Error bars represent means ± SD, one-way ANOVA followed by Tukey’s post hoc test with ***P = 0.0005 NGM vs. sorbitol, ****P < 0.0001 sorbitol vs. glucose. **g,** Expression levels of shunt genes over four hours of food deprivation followed by one hour of refeeding. Measured by qPCR, normalised to 0h time point for each gene. Error bars represent means ± SD, n = 3. **h,** DHAP levels in fed and food deprived worms (2h and 4h starvation), normalised to fed condition (n = 10). Error bars represent means ± SD, Kruskal-Wallis test followed by Dunn’s post hoc test with **P = 0.0046, ns: not significant. **i,** *gpdh-1* expression levels at control osmolarity (NGM) or 200 mM NaCl (3 hours), in the presence or absence of food (n = 3). Measured by qPCR, normalised to control treatment for each food condition. Error bars represent means ± SD, ordinary two-way ANOVA followed by Fisher’s LSD test with ****P < 0.0001, **P = 0.0027. **j,** *gpdh-1* expression levels following 30 minutes of diacetyl exposure, in the presence or absence of food (n = 6). Measured by qPCR, normalised to control treatment for each food condition. Error bars represent means ± SD, ordinary two-way ANOVA followed by Sidak’s post hoc test; *P = 0.0447 control vs diacetyl (+food), ****P < 0.0001 control vs diacetyl (-food), *P = 0.0447 +food vs -food (diacetyl).

PCA of the RNA-seq samples showed no clustering of control and diacetyl exposed groups, suggesting that diacetyl exposure did not broadly alter gene expression in our experimental model (**Fig. S1a**). Differential gene expression analysis showed no significant differences between diacetyl and control groups at 5 (**Fig. S1b**) and 15 minutes of exposure (**Fig. S1c**). The earliest significant effects appeared after 30 minutes. Twelve genes were significantly upregulated by diacetyl exposures, while no genes were downregulated (**Fig. 1b**). The upregulated genes were *gpdh-1, pqm-1, T07E3.4, ogt-1, dpy-31, gpdh-2, pgph-2, pgph-3, F20G2.1, Y51A2D.13, Y43F8B.2* and *jmjd-3.3* (**Fig. 1c**), predicted to be expressed predominantly in the intestine and alimentary system (**Fig. S1d**). Strikingly, five of these genes (*gpdh-1, gpdh-2, pgph-2, pgph-3* and *ogt-1*) are involved in the hyperosmotic stress response through their role in glycerol production. GPDH-1 and GPDH-2 convert DHAP into Gro3P, which is converted by PGPH-2 and PGPH-3 into glycerol. Together, these two reactions form part of the DHAP-Glycerol shunt (**Fig. 1d**). *ogt-1*, a conserved O-GlcNAc transferase, was recently described as an essential regulator of the hyperosmotic stress response that enables *gpdh-1* upregulation^27^.

In addition to their canonical role in the hyperosmotic stress response, enzymes in the DHAP-Glycerol shunt have recently emerged as regulators of metabolic stress and glucotoxicity across organisms^16, 18^. Given their prevalence in the gene expression response to a food cue, we aimed to better characterize their regulation in response to food availability and glucose. A previous study^17^ observed *gpdh-1* upregulation in worms raised on a high glucose diet (NGM supplemented with 2% glucose), but did not clarify whether this upregulation was caused by the metabolic or the osmotic effects of glucose supplementation. We observed that plates supplemented with 2% glucose induced a *gpdh-1* transcriptional reporter to a greater extent than plates supplemented with sorbitol at the same concentration (111 mM), suggesting that excess glucose induced *gpdh-1* by causing both an osmotic and a metabolic stress (**Fig. 1e - f**). We hypothesised that the DHAP-Glycerol shunt may also be regulated in response to feeding. To test this, we measured expression of shunt genes during food deprivation and refeeding (**Fig. 1g**). *gpdh-1*, the gene most upregulated by diacetyl exposure, was rapidly repressed at the onset of food deprivation and upregulated after refeeding. Of the other three genes measured, only *pgph-2* responded similarly to *gpdh-1*, although to a lesser extent. Beyond this short fasting timeframe, we showed that *gpdh-1* remained lowly expressed throughout 24 hours of food deprivation (**Fig. S1e**), confirming that the pathway is suppressed by starvation, potentially to maintain DHAP and GAP levels. Through our metabolomic analysis, we observed that along with glucose itself and GAP, DHAP was in fact one of the glycolytic intermediates significantly depleted at the onset of food deprivation (**Fig. 1h** and **S1f**).

Knowing that *gpdh-1* expression is induced by hyperosmotic stress and repressed by food deprivation, we investigated how these opposing signals would interact when combined. Under conditions of food deprivation, hyperosmotic stress failed to upregulate expression of the *gpdh-1* reporter to the levels observed in fed conditions (**Fig. S1g**). Using qPCR for a more sensitive readout, we confirmed that food deprivation reduced *gpdh-1* induction under hypertonic conditions (**Fig. 1i** and **S1h**), suggesting that food availability is the dominant input governing the expression of this gene (**Fig. 1i**). Having established this, we next wanted to test whether diacetyl exposure may be acting as a hyperosmotic stressor or as a food signal. Similarly to above, we dissected the overlap of food deprivation and diacetyl signals by exposing worms to diacetyl in the presence and absence of food. We observed that unlike hyperosmotic stress, diacetyl exposure upregulated *gpdh-1* expression more strongly during food deprivation: a 5.8 fold in the absence of food against a 2.4 fold increase in the presence of food (**Fig. 1j**). Together, these results show that *gpdh-1* expression, induced by diacetyl and hyperosmotic stress, is also more broadly induced by nutrient availability in the form of excess glucose or food. We hypothesise that diacetyl may act as a preemptive food signal that partially overlaps with the response to food itself.

### Diacetyl triggers metabolic shift overlapping with hyperosmotic stress response

Having observed a rapid transcriptional upregulation of enzymes in worms exposed to diacetyl, we wanted to assess how this food cue would alter the metabolic profile of fasting worms. Using metabolomics and lipidomics, we measured the effects of volatile diacetyl exposure in the same experimental model used for transcriptomics. We found that one hour of diacetyl exposure induced a metabolic shift in glycerol containing phospholipids, matching the transcriptional signature observed after 30 minutes of diacetyl exposure. Diacetyl exposure altered the levels of 134 metabolites (77 increased, 57 decreased), 80 of these being glycerophospholipids (**Supplementary Table 1**). Diacetyl specifically increased the levels of 14 out of the 21 detected phosphatidylglycerols (PGs) (**Fig. 2a**), particularly PG(16:0_20:2), PG(16:0_18:1) and PG(16:0_20:1), all containing palmitic acid (16:0). PGs are glycerophospholipids with a glycerol head group, synthesised by a pathway that consumes phosphatidic acid (PA), Gro3P and CTP (**Fig. 2b**). Accordingly, diacetyl exposure decreased levels of 9 out of the 18 detected PA species (**Fig. S2a**), except PA(18:2_18:2) which was increased. Diacetyl did not change levels of DHAP and Gro3P, suggesting that these metabolites may be replenished after being consumed for PG synthesis.

**Figure 2.**
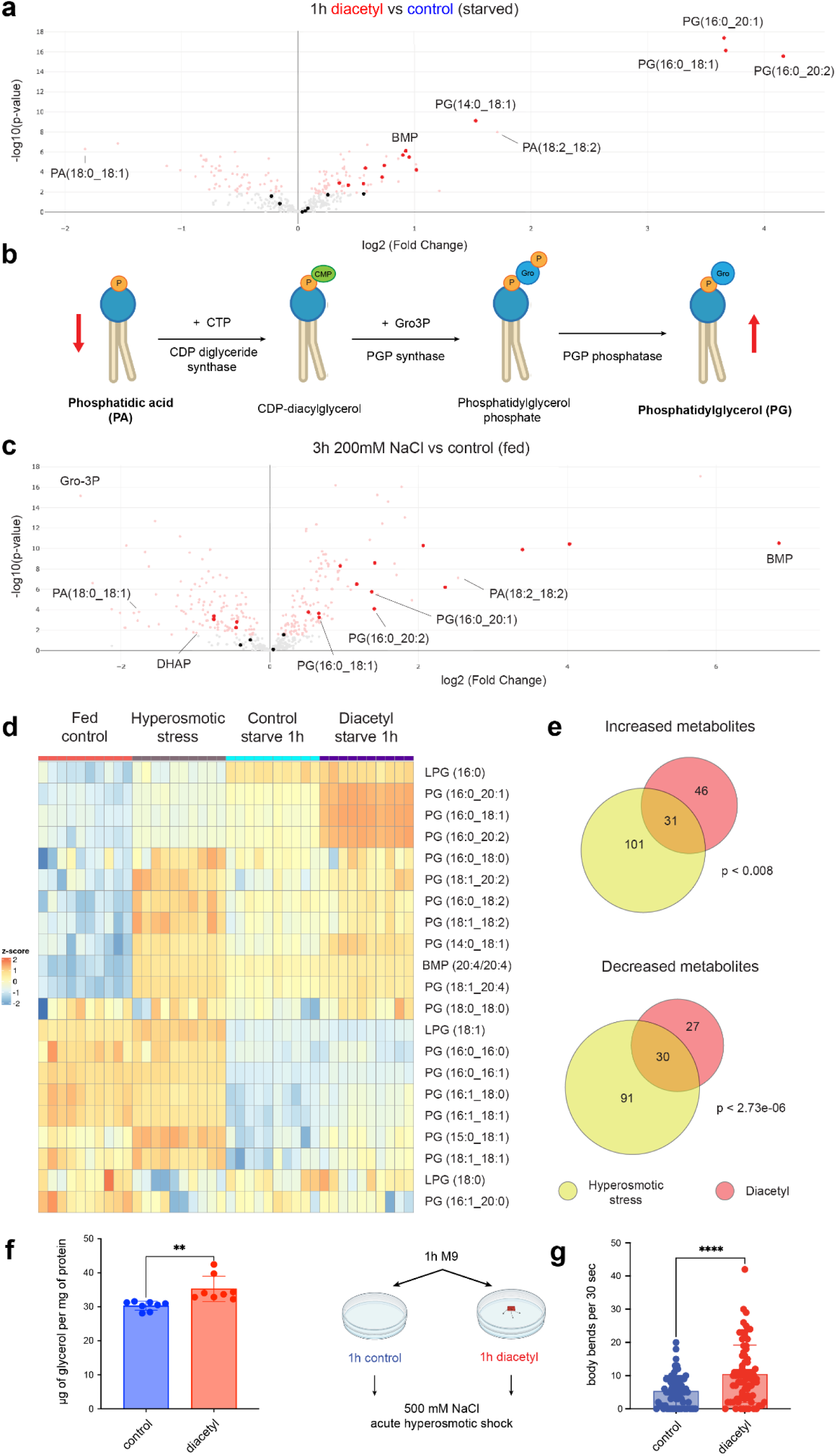
Diacetyl triggers metabolic shift overlapping with hyperosmotic stress response. **a**, Volcano plot comparing metabolomic effects of diacetyl exposure (one hour) to control food deprivation. Metabolites significantly altered in red (FDR <0.05, Welch’s t-test). Highlighted are metabolites classified as phosphatidylglycerols (PGs). **b,** Schematic depicting synthesis of PGs from phosphatidic acid (PA) precursor lipids and Gro3P. **c,** Volcano plot comparing metabolomic effects of hyperosmotic stress (three hours, 200 mM NaCl plates with food) to control NGM plates. Metabolites significantly altered in red (FDR <0.05, Welch’s t-test). Highlighted are metabolites classified as PGs. **d,** Heatmap comparing levels of all phosphatidylglycerols (PGs) in fed condition, hyperosmotic stress (three hours, 200 mM NaCl), one hour food deprivation (control) and one hour diacetyl exposure during food deprivation (n = 10). **e,** Overlap between metabolites significantly altered (FDR <0.05, Welch’s t-test) by hyperosmotic stress (three hours, 200 mM NaCl plates with food) and diacetyl exposure (one hour, in the absence of food). Significance determined by hypergeometric test; increased metabolites x = 31, N = 472, p < 0.008; decreased metabolites x = 30, N = 472, p < 2.73e-06. **f,** Worm glycerol levels following two hours of diacetyl exposure, normalised by protein content. Error bars represent means ± SD, Welch’s t-test with **P = 0.0065; n =8. **g,** Effects of diacetyl pre-exposure on acute hyperosmotic stress resistance. Worms were pre-exposed to diacetyl for one hour (in the absence of food), before thrashing was recorded on hyperosmotic stress plates (500 mM NaCl). Data is pooled from three biological replicates; n = 71 worms in the control group; n = 78 in the diacetyl group. Error bars represent means ± SD, Mann-Whitney test with ****P < 0.0001.

To test whether these metabolic signatures are linked to DHAP-Glycerol shunt upregulation, we compared the effects of diacetyl exposure to the effects of hyperosmotic stress, which also induces this pathway^23^. We exposed worms (in the presence of food) to 200 mM NaCl for three hours. Hyperosmotic stress increased the levels of 13 out of the 21 detected PGs (**Fig. 2c**) and decreased levels of 10 out of the 18 detected PAs (except PA(18:2_18:2)) (**Fig. S2b**). Significantly, a similar set of PG species were increased by diacetyl exposure and hyperosmotic stress (**Fig. 2d**). While hyperosmotic stress induced broader metabolic changes (132 metabolites increased, 121 decreased) (**Supplementary Table 1**) compared to the food cue, the nature and direction of the changes resembled the diacetyl effects. In fact, we observed a significant overlap between all the metabolites altered by these two stimuli (**Fig. 2e**). Unlike diacetyl, hyperosmotic stress reduced levels of both DHAP and Gro3P, the latter being the metabolite most downregulated by this stress. While the metabolomic approach we employed did not detect glycerol, using a colorimetric kit, we observed that diacetyl exposure led to a significant increase in whole-worm glycerol levels (**Fig. 2f**).

Given the overlap in metabolic responses between the volatile exposure and hyperosmotic stress, we tested whether diacetyl exposure and the subsequent metabolic remodelling we observed protected worms against hyperosmotic stress. We assessed this by measuring the thrashing rate of worms placed on high NaCl plates. We found that one hour pre-exposure to diacetyl strongly protected worms from this acute hyperosmotic stress (**Fig. 2g**), without affecting thrashing rates in unstressed conditions (**Fig. S2c**). These results point to an overlap in diacetyl and hyperosmotic responses not only at the transcriptional and metabolic levels, but also in stress adaptation.

### Prolonged diacetyl exposure in the absence of food boosts FMO-2 dependent responses

While DHAP-Glycerol shunt genes were no longer differentially expressed by 90 minutes, our transcriptomic time course (**Fig. 3a**) revealed upregulation of a second wave of gene expression changes. At this time point, nine genes were significantly upregulated by diacetyl, while no genes were downregulated (**Fig. 3b**). The upregulated genes were *C45E5.1, C06E4.3, F56D5.3, fmo-2, D1054.8, F20G2.1, cth-1, argk-1* and *eef-1A.2* (**Fig. 3c**). This wave was characterised by a strong upregulation of *fmo-2*, a flavin-containing monooxygenase that is a well-established marker of the dietary restriction (DR) response, as well as being required for the longevity effects of DR^25^.

**Figure 3.**
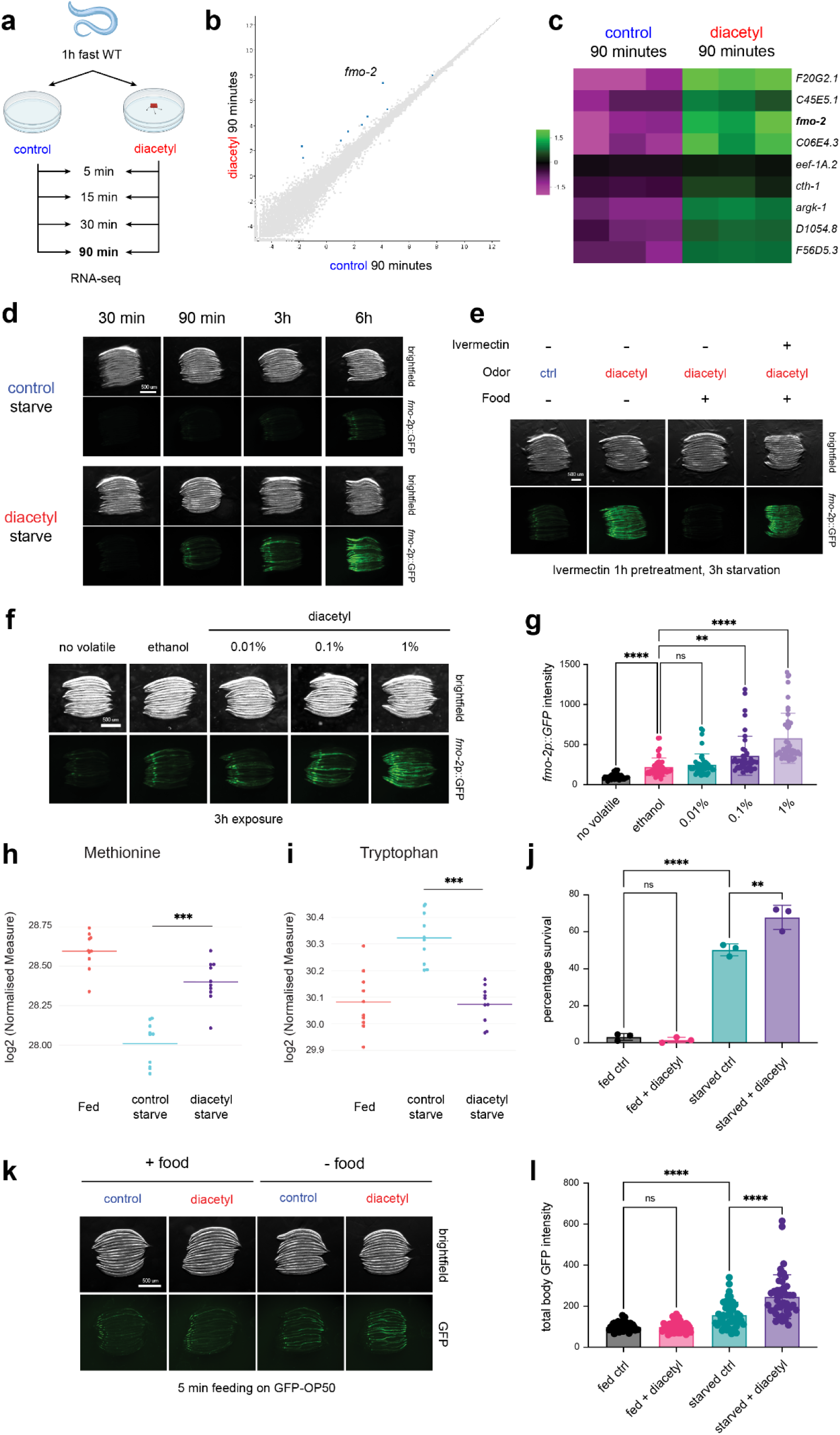
Prolonged diacetyl exposure in the absence of food boosts FMO-2 dependent responses. **a**, Experimental setup for measuring gene expression effects of diacetyl exposure in food-deprived worms, comparing control food deprivation (no volatile) to food deprivation in the presence of volatile diacetyl (1% solution in ethanol, pipetted on plate lid). Three biological replicates collected for RNA-seq over a ninety-minute time course. **b,** Scatter plot showing differential gene expression between control and diacetyl groups at 90 minutes of exposure. **c,** Heatmap showing differential gene expression between control and diacetyl groups at 90 minutes of exposure. **d,** Representative images of *fmo-2* transcriptional reporter induction over six hours of food deprivation, in the presence or absence of diacetyl. Images shown are from one experiment out of three biological replicates. Additional replicates are available in Figure 3 - Source data d. **e,** Representative images of *fmo-2* reporter worms treated with ivermectin for one hour, followed by diacetyl exposure in the presence or absence of food for three hours. Images shown are from one experiment out of three biological replicates. Additional replicates are available in Figure 3 - Source data e. **f,** Representative images of *fmo-2* reporter worms exposed to no volatile (control), ethanol or diacetyl dilutions in ethanol, over three hours of food deprivation. Images shown are from one experiment out of three biological replicates. Additional replicates are available in Figure 3 - Source data f. **g,** Quantification of reporter induction in experiment from (f). Data is pooled from all three replicates (n = 45 worms per condition). Error bars represent means ± SD, Brown-Forsythe and Welch ANOVA test followed by Dunnett’s T3 post hoc test, ****P < 0.0001 vs no volatile, **P = 0.0035 vs 0.1% diacetyl, ****P < 0.0001 vs 1% diacetyl, ns: not significant. **h** - **i,** Individual metabolite levels in fed and food deprived worms, starved either in control conditions or during diacetyl exposure for one hour (n = 10). Log2 normalised levels on vertical axes. **h,** Methionine levels, Welch’s t-test with ***P = 0.0001. **i,** Tryptophan levels, Welch’s t-test with ***P = 0.0001. **j,** Survival (day 3 of adulthood) of worms exposed for three hours to diacetyl in the presence or absence of food (day 1) and heat-shocked on day 2. Error bars represent means ± SD (n = 3), ordinary one-way ANOVA followed by Sidak’s post hoc test with ****P < 0.0001 fed vs starved (ctrl), **P = 0.0015 ctrl vs diacetyl (starved), ns: not significant. **k,** Representative images of worms fed with GFP expressing bacteria following a three-hour exposure to diacetyl in the presence or absence of food. Images shown are from one experiment out of three biological replicates. Additional replicates are available in Figure 3 - Source data k. **l,** Quantification of GFP intensity in experiment from (k). Data is pooled from all three replicates (n = 45 worms per condition). Error bars represent means ± SD, Brown-Forsythe and Welch ANOVA test followed by Dunnett’s T3 post hoc test, ****P < 0.0001, ns: not significant.

Using a transcriptional *fmo-2* reporter we confirmed that diacetyl exposure dramatically accelerated the induction of *fmo-2* during food deprivation (**Fig. 3d**). This effect began as early as thirty minutes of exposure (**Fig. S3a**), and stabilised around ninety minutes, resulting in a four-fold induction of *fmo-2* with diacetyl exposure compared to control starvation (**Fig. S3b**). Critically, we observed that this diacetyl effect was dependent on food deprivation, as exposing worms to diacetyl in the presence of food (OP50) suppressed *fmo-2* induction (**Fig. 3e**). We wondered whether this OP50 suppressive effect depended on the worms ingesting their food or simply on conflicting sensory signals coming from the bacteria. To test this, we briefly pretreated worms with ivermectin, blocking pharyngeal pumping, before exposure to diacetyl. We found that ivermectin completely reversed the OP50 suppressive effect, showing that food ingestion is the mechanism by which OP50 suppresses the *fmo-2* diacetyl response (**Fig. 3e**). We also confirmed that ivermectin treatment by itself did not induce *fmo-2* (**Fig. S3c**).

Diacetyl has been shown to be chemoattractive to worms at dilutions as low as 0.0001%. Some studies have found aversive diacetyl responses at concentrations of 10%^28^ or 100%^29^. We decided to test the relationship between *fmo-2* expression levels and diacetyl over a range of non-aversive dilutions. We exposed worms to ten-fold dilutions of diacetyl and found that this induced *fmo-2* more than the ethanol control in the 0.1% to 1% dilution range (**Fig. 3f - g**). Interestingly, the ethanol control itself induced *fmo-2* when compared to the worms exposed to no volatiles (**Fig. 3f - g**). To investigate the sensory mechanisms mediating diacetyl gene expression effects, we first tested two diacetyl receptor mutants. ODR-10 mediates diacetyl chemotaxis in the 0.1% to 1% dilution range^30^, while SRI-14 mediates aversion to undiluted diacetyl^31^. We observed that both *odr-10* and *sri-14* mutants induced *fmo-2* normally in response to our standard 1% diacetyl treatment (**Fig. S3d**). Looking more broadly at signalling pathways involved in food sensing^32^, we found that mutants in serotonin and dopamine signalling also remained responsive to diacetyl (**Fig. S3e**). Likewise, mutations impairing neurotransmitter or neuropeptide release (*unc-13* and *unc-31* respectively) did not suppress *fmo-2* induction by diacetyl (**Fig. S3f**), hinting at a potential non-neuronal sensing mechanism. To test this, we used a *daf-19* (*m86*); *daf-12* (*sa204*) strain completely lacking all sensory cilia^33^. Surprisingly, even these worms responded to diacetyl exposure by strongly expressing *fmo-2* (**Fig. S3g**). Together, these results suggest that the acute gene expression effects we observe in our study do not depend on canonical diacetyl sensing mechanisms or ciliated olfactory neurons.

We then explored the effects of prolonged diacetyl exposure beyond gene expression, linking FMO-2 upregulation to biochemical, physiological and behavioural readouts. Higher *fmo-2* expression levels have been shown to correlate with higher levels of methionine and decrease levels of tryptophan, a direct substrate of FMO-2^26^. Diacetyl exposure increased levels of methionine (**Fig. 3h**) and decreased levels of tryptophan (**Fig. 3i**), consistent with an increase in FMO-2 activity in this condition. Because *fmo-2* overexpression boosts heat shock resistance^25^, we decided to test if diacetyl exposure could also improve this phenotype. We found that a three-hour starvation period on day 1 greatly improved thermotolerance on day 2, as assessed by thermorecovery on day 3 (**Fig. 3j**). Diacetyl exposure combined with this starvation treatment further protected the worms, while diacetyl exposure on food had no effect (**Fig. 3j**). Notably, this effect was *fmo-2* dependent (**Fig. S3h**). Finally, FMO-2 has been recently described to regulate foraging behaviours and satiety responses^34^. Food deprivation is also known to increase feeding rates (appetite) in *C. elegans*^35^. We found that diacetyl exposure in the absence of food increased appetite upon refeeding, while exposure to diacetyl on food did not affect feeding rates (**Fig. 3k** - **l**). Again, we observed that this effect was *fmo-2* dependent (**Fig. S3i**).

Together, these results indicate that prolonged diacetyl exposure during food deprivation accelerates the induction of *fmo-2* through non-canonical sensing mechanisms, and boosts FMO-2 related phenotypes.

### Diacetyl and hyperosmotic stress induce *fmo-2* through DHAP-Glycerol shunt induction

The temporal signature of DHAP-Glycerol shunt induction (starting from 30 minutes of diacetyl exposure), closely followed by *fmo-2* upregulation (peaking at 90 minutes of exposure), suggested a potential causal relationship between these two responses (**Fig. 4a**). To test this, we knocked down shunt components and assayed *fmo-2* expression. We observed that RNAi knockdown of the shunt components *pgph-2* and *pgph-3* significantly reduced the diacetyl-induced expression of *fmo-2*.(**Fig. 4b - c**), establishing a causal link between the DHAP-Glycerol shunt and FMO-2.

**Figure 4.**
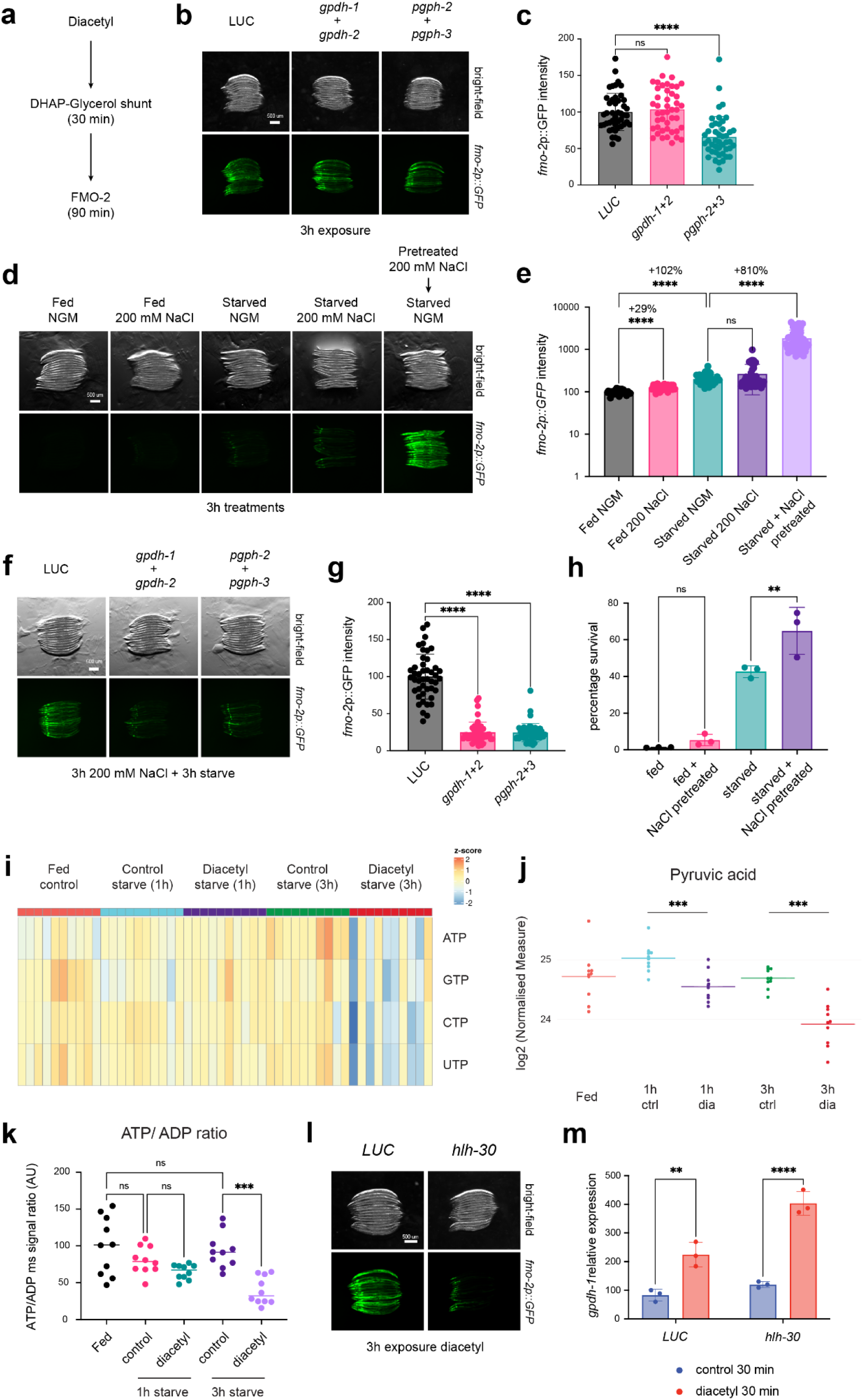
Diacetyl and hyperosmotic stress induce *fmo-2* through DHAP-Glycerol shunt induction. **a**, Schematic of gene expression responses to diacetyl exposure, detected from transcriptomics time course. **b,** Representative images of *fmo-2* transcriptional reporter induction by diacetyl exposure in the absence of food (three hours), in worms raised on non-targeting Luciferase (LUC), combined RNAi targeting DHAP shunt components *gpdh-1* and *gpdh-2* or *pgph-2* and *pgph-3*. Images shown are from one experiment out of three biological replicates. Additional replicates are available in Figure 4 - Source data b. **c,** Quantification of reporter induction in experiment from (b). Data is pooled from all three replicates (n = 45 worms per condition). Error bars represent means ± SD, Ordinary one-way ANOVA followed by Dunnett’s T3 post hoc test with ****P < 0.0001, ns: not significant. **d,** Representative images of *fmo-2* transcriptional reporter induction by three hours of hyperosmotic stress (Fed, 200 mM NaCl), food deprivation (Starved, NGM), hyperosmotic stress combined with food deprivation (Starved, 200 mM NaCl) and hyperosmotic stress pretreatment in the presence of food followed by food deprivation at control osmolarity (Pretreated 200 mM NaCl + Starved NGM). Images shown are from one experiment out of three biological replicates. Additional replicates are available in Figure 4 - Source data d. **e,** Quantification of reporter induction in experiment from (d). Data is pooled from all three replicates (n = 45 worms per condition). Log 10 y-axis. Error bars represent means ± SD, Brown-Forsythe and Welch ANOVA tests followed by Dunnett’s T3 post hoc test with ****P < 0.0001, ns: not significant. **f,** Representative images of *fmo-2* transcriptional reporter induction by hyperosmotic stress pretreatment (200 mM NaCl) in the presence of food followed by food deprivation at control osmolarity, in worms raised on non-targeting Luciferase (LUC), combined RNAi targeting DHAP shunt components *gpdh-1* and *gpdh-2* or *pgph-2* and *pgph-3*. Images shown are from one experiment out of three biological replicates. Additional replicates are available in Figure 4 - Source data f. **g,** Quantification of reporter induction in experiment from (f). Data is pooled from all three replicates (n = 45 worms per condition). Error bars represent means ± SD, Brown-Forsythe and Welch ANOVA tests followed by Dunnett’s T3 post hoc test with ****P < 0.0001. **h,** Survival (day 3 of adulthood) of worms exposed on day 1 to 200 mM NaCl in the presence of food, followed by three hours at control osmolarity in the presence (fed + NaCl pretreated) or absence of food (starved + NaCl pretreated). Worms heat-shocked on day two. Error bars represent means ± SD (n = 3), ordinary one-way ANOVA followed by Sidak’s post hoc test with **P = 0.0077, ns: not significant. **i,** Heatmap comparing levels of ribonucleoside triphosphates in fed conditions and food deprivation (Starve, one or three hours) in the presence or absence (Control) of diacetyl exposure (n = 10). **j,** Pyruvic acid levels across samples in fed worms, starved worms in control conditions (ctrl), starved worms exposed to diacetyl (1 and 3 hour timepoints) (n = 10). Log2 normalised levels on vertical axes. Welch’s t-test with ***P = 0.0003 (1h) and ***P = 0.0009 (3h). **k,** ATP to ADP ratios of LC-MS peak area measurements (Arbitrary Units) detected across samples in fed worms, starved worms in control conditions, starved worms exposed to diacetyl (1 and 3 hour timepoints) (n = 10). Brown-Forsythe and Welch test followed by Dunnett’s T3 post hoc tests with ***P = 0.0001. **l,** Representative images of *fmo-2* transcriptional reporter induction by diacetyl exposure in the absence of food (three hours), in worms raised on non-targeting Luciferase (LUC) or RNAi targeting *hlh-30*. Images shown are from one experiment out of three biological replicates. Additional replicates are available in Figure 4 - Source data l. **m,** *gpdh-1* expression levels following 30 minutes of diacetyl exposure, in worms raised on non-targeting Luciferase (LUC) or RNAi targeting *hlh-30*. Measured by qPCR, normalised to LUC control. Error bars represent means ± SD (n = 3), ordinary two-way ANOVA followed by Sidak’s post hoc test; **P = 0.0013 (LUC), ****P < 0.0001 (hlh-30).

Next, to test if shunt activation may be sufficient to induce *fmo-2* in the absence of a food cue, we leveraged its induction by hyperosmotic stress. Hyperosmotic stress in the presence of food moderately induced (29% increase) *fmo-2* expression. In comparison, three hours of starvation doubled *fmo-2* expression (**Fig. 4d - e**). Pairing hyperosmotic stress and starvation did not increase FMO-2 induction. Having previously shown that food deprivation suppresses shunt induction (**Fig. 1i** and **S1h**), we pretreated worms with 200 mM NaCl while on food, before starting starvation at regular osmolarity. We found that this pretreatment induced *fmo-2* by eight-fold (**Fig. 4d - e**), an effect even stronger than diacetyl exposure. This induction could be suppressed by combined RNAi targeting either *gpdh-1* and *gpdh-2* or *pgph-2* and *pgph-3* (**Fig. 4f - g**), demonstrating that the DHAP-Glycerol shunt is the critical factor inducing *fmo-2* regardless of the upstream trigger. We also tested if this effect on *fmo-2* expression would be coupled with a boost in starvation-related benefits. Indeed, we found that pairing the 200 mM NaCl pretreatment with food deprivation increased the beneficial effects on heat shock recovery (**Fig. 4h**), an effect not seen with hyperosmotic treatment on its own, mimicking the effects of diacetyl on thermotolerance.

To identify potential metabolic triggers of *fmo-2* expression, we compared metabolomic signatures of diacetyl after one and three hours of exposure. The two timepoints strongly overlapped in metabolic responses to diacetyl (**Fig. S4a**). After three hours, diacetyl exposed worms still showed higher levels of PGs, including the three species most accumulated starting from one hour of exposure (**Fig. S4b - c**). In addition, after three hours of exposure diacetyl depleted the cellular ribonucleoside triphosphate (NTP) pool (**Fig. 4i**), significantly decreasing levels of CTP, UTP and ATP (**Fig. S4d - f**). CTP is the precursor used to produce CDP-diacylglycerol (**Fig. 2b**), and its levels are replenished by CTP synthase, using UTP as a substrate and consuming ATP in the reaction^36^. We hypothesized that inducing the DHAP-Glycerol shunt during food deprivation may force an anabolic reaction diverting glycolytic intermediates and consuming cellular NTPs in the process. In fact, diacetyl exposure also reduced levels of pyruvic acid starting from one hour of exposure (**Fig. 4j**) and strongly decreased the ATP/ADP ratio (**Fig. 4k**) at three hours of exposure, matching the *fmo-2* induction at this timepoint.

We then investigated transcription factors potentially linking DHAP-Glycerol shunt induction and metabolic remodelling to *fmo-2* expression. Low ATP levels during starvation activate AMPK to induce TFEB-dependent transcriptional programs^37^. We observed that knocking down *hlh-30* by RNAi completely suppressed *fmo-2* induction by diacetyl (**Fig. 4l**). *hlh-30* RNAi did not however block *gpdh-1* induction by diacetyl (**Fig. 4m**), nor did it block the diacetyl mediated hyperosmotic stress protection (**Fig. S4g**), reinforcing the hypothesis that it acts downstream or in parallel to shunt induction.

These findings show that the induction of the DHAP-Glycerol shunt precedes and is sufficient for *fmo-2* induction in the absence of food, regardless of whether the specific trigger is diacetyl or hyperosmotic stress. Prolonged diacetyl exposure is associated with depleted levels of high-energy phosphate donors and ATP to ADP ratios, potentially leading to *fmo-2* expression through HLH-30 activation.

### *gpdh-1* induction and metabolic effects of diacetyl depend on MDT-15

Having established that diacetyl exposure induced a transcriptional response related to glucose excess, we wondered whether regulators of glucotoxicity might also mediate diacetyl transcriptional responses. MDT-15 and NHR-49 are key transcription factors modulating lipid metabolism^38^ and glucotoxicity resistance^39^. Both genes have also been previously linked to the starvation response^40, 41^ and *fmo-2* expression^42^. Similarly to HLH-30, we found that their knockdown completely suppressed *fmo-2* induction by diacetyl (**Fig. S5a**). Unlike HLH-30 however, both *nhr-49* and *mdt-15* RNAi significantly suppressed *gpdh-1* induction by diacetyl (**Fig. 5a**), suggesting they act upstream of DHAP-Glycerol shunt induction. Additionally, *mdt-15* knockdown completely blocked the protective effect of diacetyl exposure on acute hyperosmotic stress resistance (**Fig. 5b**).

**Figure 5.**
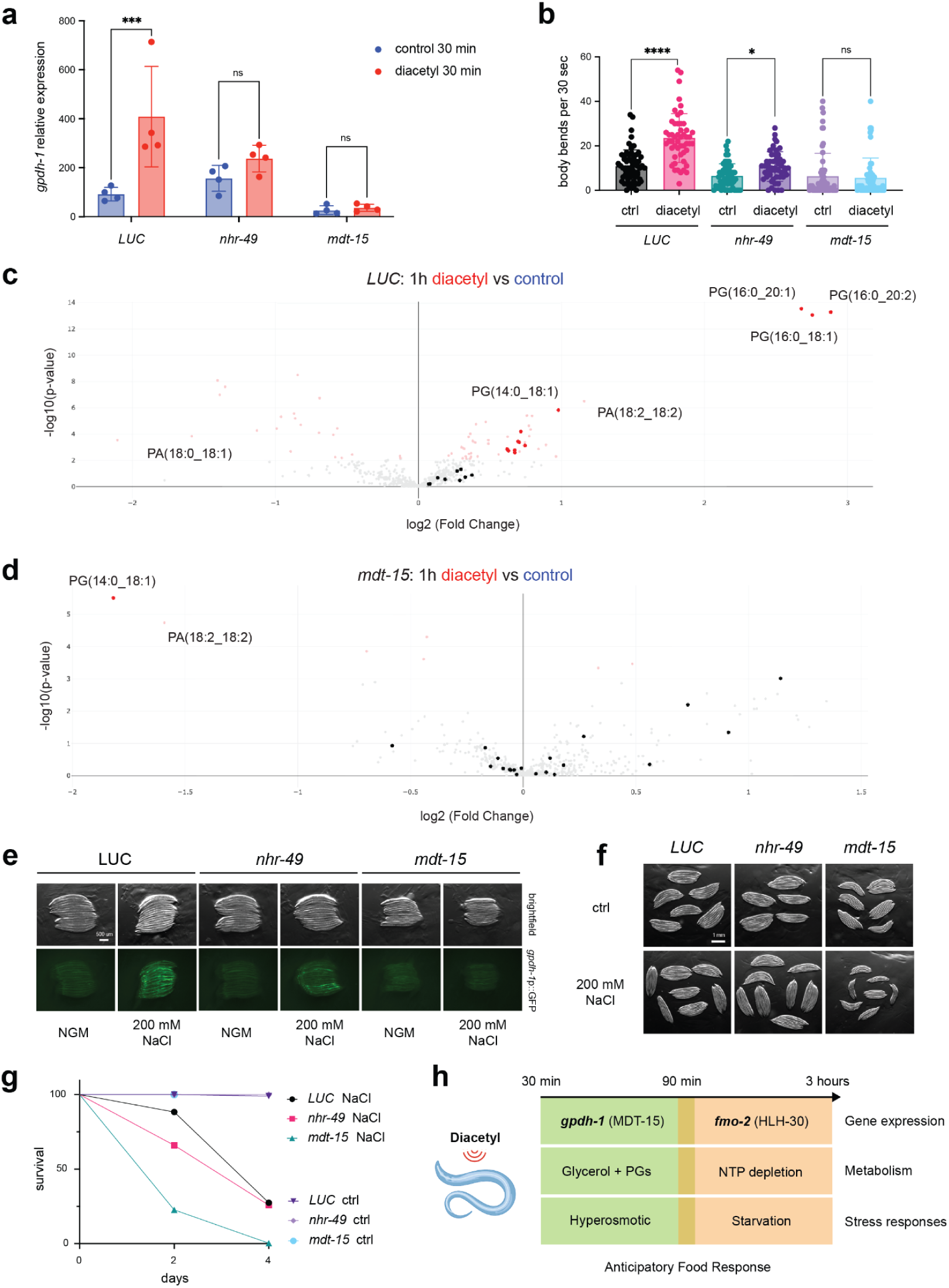
*gpdh-1* induction and metabolic effects of diacetyl depend on MDT-15. **a**, *gpdh-1* expression levels following 30 minutes of diacetyl exposure, in worms raised on non-targeting Luciferase (LUC) RNAi targeting *nhr-49* or *mdt-15*. Measured by qPCR, normalised to LUC control. Error bars represent means ± SD, ordinary two-way ANOVA followed by Sidak’s post hoc test with ***P = 0.0003, ns: not significant. n = 4. **b**, Effects of diacetyl pre-exposure on acute hyperosmotic stress resistance. Worms raised on non-targeting Luciferase (LUC) RNAi targeting *nhr-49* or *mdt-15* were pre-exposed to diacetyl for one hour (in the absence of food), before thrashing was recorded on hyperosmotic stress plates (500 mM NaCl). Data is pooled from three biological replicates; n = 64 worms (LUC, ctrl), n = 51 (LUC, diacetyl), n = 55 (*nhr-49*, ctrl), n = 50 (*nhr-49*, diacetyl), n = 60 (*mdt-15*, ctrl), n = 55 (*mdt-15*, diacetyl). Error bars represent means ± SD, ordinary one-way ANOVA followed by Sidak’s post hoc test with ****P < 0.0001 (LUC), *P = 0.03 (*nhr-49*), ns: not significant. **c** - **d,** Volcano plots comparing metabolomic effects of diacetyl exposure (one hour) to control food deprivation, for worms raised on Luciferase (LUC) RNAi (**c**) or *mdt-15* RNAi (**d**). Metabolites significantly altered in red (FDR <0.05, Welch’s t-test). Highlighted are metabolites classified as phosphatidylglycerols (PGs). **e**, Representative images of *gpdh-1* transcriptional reporter worms raised on Luciferase (LUC), *nhr-49* and *mdt-15* targeting RNAi and exposed to 200 mM NaCl for three hours. Images shown are from one experiment out of three biological replicates. Additional replicates are available in Figure 5 - Source data e. **f**, Representative images of worms raised on Luciferase (LUC), *nhr-49* and *mdt-15* targeting RNAi, either at control osmolarity or in the presence of 200 mM NaCl. Images shown are from one experiment out of three biological replicates. Additional replicates are available in Figure 5 - Source data f. **g**, Representative survival of worms raised on Luciferase (LUC), *nhr-49* and *mdt-15* targeting RNAi and transferred on day 1 of adulthood to RNAi plates supplemented with 400 mM NaCl. Results shown are from one experiment out of three biological replicates. Additional replicates are available in Figure 5 - Source data g. **h**, Model summarising acute effects of volatile diacetyl exposure across the levels of gene expression, metabolism and physiological stress responses. Food deprived worms exposed to diacetyl induce expression of *gpdh-1* (through MDT-15) and other enzymes in the DHAP-glycerol shunt. This leads to accumulation of glycerol, PGs and protection from acute hyperosmotic stress. The metabolic shift induced by the shunt, in combination with food deprivation (possibly sensed through HLH-30), then triggers a state of enhanced starvation characterised by depleted NTP levels, upregulated *fmo-2* expression and starvation-related phenotypes.

We focused on *mdt-15* RNAi, which showed the strongest effect on *gpdh-1* suppression, to test if blocking shunt induction by diacetyl would also reverse the metabolic effects of diacetyl exposure. To do this, we compared the diacetyl metabolic response of worms grown either on Luciferase (LUC) non-targeting RNAi or *mdt-15* RNAi. In the LUC RNAi condition, one hour of diacetyl exposure largely recapitulated the effects measured in WT worms (**Fig. 5c**): it increased the levels of PGs, particularly PG(16:0_20:2), PG(16:0_18:1) and PG(16:0_20:1), while decreasing levels of PAs (**Fig. S5b**), except PA(18:2_18:2). *mdt-15* RNAi completely abolished these effects (**Fig. 5d**). In these worms, diacetyl exposure altered the abundance of only 7 metabolites, compared to the 70 in LUC controls. No PGs were increased by diacetyl. PG(14:0_18:1), one of the metabolites most induced by diacetyl in WT and LUC ctrl worms, was in fact decreased by diacetyl in *mdt-15* RNAi. Likewise PA(18:2_18:2), induced by diacetyl in WT and LUC control worms, was decreased in *mdt-15* RNAi. We conclude that MDT-15 is required for the metabolic remodeling downstream of *gpdh-1* induction by diacetyl.

We next asked if NHR-49 and MDT-15 could regulate shunt induction independently of nutritional stressors. Surprisingly, we found that *nhr-49* knockdown reduced *gpdh-1* induction by hyperosmotic stress (200 mM NaCl), and *mdt-15* knockdown completely suppressed it (**Fig. 5e** and **S5c**). Accordingly, we tested if *mdt-15* and *nhr-49* would also be required for responses to hyperosmotic stress, finding that knockdown of *mdt-15* impaired development on 200 mM NaCl (**Fig. 5f** and **S5d**). Knockdown of *mdt-15* also reduced survival over 4 days of hyperosmotic stress treatment with 400 mM NaCl (**Fig. 5g**), without affecting survival over this time-window in control conditions. Knockdown of *nhr-49* did not significantly affect development or survival on hyperosmotic stress. These results demonstrate that MDT-15 is required for *gpdh-1* induction by diacetyl and hyperosmotic stress, as well as for the diacetyl induced metabolic remodelling and hyperosmotic stress resistance (**Fig. 5h**).

## Discussion

In this study, we characterized the acute responses of fasting *C. elegans* exposed to a food-related volatile cue. We found that exposure to diacetyl induces interconnected transcriptional and metabolic responses that unfold across two temporal phases (**Fig. 5h**). The first phase is initiated by the rapid expression of *gpdh-1* and other enzymes comprising the DHAP-Glycerol shunt. The second is defined by the subsequent upregulation of the metabolic regulator FMO-2.

Previous work has shown that enzymes in the DHAP-Glycerol shunt are upregulated to produce glycerol during hyperosmotic stress^20,22^, and to protect worms against glucotoxicity^20^. Going beyond this, we demonstrate that this pathway is also highly responsive to food availability (**Fig. 1g**) and to a food cue, underscoring its role in regulating nutrient homeostasis. While DHAP and GAP have been reported as the glycolytic intermediates most accumulated during glucotoxicity^14^, we show that they are also depleted during food deprivation (**Fig. 1h** and **S1f**). In these conditions, the shunt may be actively suppressed to maintain efficient glycolytic flux and pyruvate synthesis, a regulatory principle similarly observed in yeast where the ortholog GPD2 is inhibited under limiting nutrients^43^. We propose that fasting worms sensing diacetyl may preemptively induce the DHAP-Glycerol shunt in preparation for food ingestion. We speculate that diacetyl, produced by LAB to dispose of excess pyruvate^7^, may be acting as a signal for the presence of pyruvate-rich bacteria. In agreement with this concept, we find that diacetyl exposure decreases pyruvate levels in the worms (**Fig. 4j**). This expands the concept of worms utilising volatile cues to gain information about the specific nutritional state of food in their environment^44^.

Following transcriptional induction of the shunt, diacetyl exposure drives a rapid metabolic remodeling event characterized by the accumulation of glycerol and phosphatidylglycerols (PGs), a metabolic shift that overlaps with the effects of hyperosmotic stress (**Fig. 2**). PGs are an understudied lipid species in *C. elegans*. They are synthesized in mitochondria, processed in the ER, and transported back to mitochondria^45^. Within mitochondria, PGs serve as precursors for the synthesis of cardiolipin, an essential component of the inner mitochondrial membrane that supports electron transport chain function^46^. While PG accumulation after diacetyl exposure may be a simple byproduct of increased Gro3P availability (stemming from GPDH-1 activity), it will be interesting to study whether diacetyl-induced PG accumulation directly affects cellular respiration or other mitochondrial functions. We identified MDT-15 as the critical regulator controlling both the diacetyl-induced *gpdh-1* expression (**Fig. 5a**) and downstream metabolic remodeling (**Fig. 5d**). Interestingly, MDT-15 appears to be broadly required for the induction of the DHAP-Glycerol pathway, including its activation under hyperosmotic stress (**Fig. 5e**). This expands the known functions of MDT-15 in mediating the expression of fatty acid desaturases during fasting^41^ and glucotoxicity^39^. This novel regulatory role over the DHAP-Glycerol shunt may also contribute to MDT-15’s protective effects during glucotoxicity, aligning with previous observations pointing to DHAP as a candidate toxic molecule mediating the lifespan shortening effects of excess glucose^39^.

The secondary phase of the diacetyl response is marked by a strong upregulation of FMO-2, a metabolic regulator implicated in the longevity effects of DR^25^, as well as in responses to hypoxia and oxidative stress^42^. Diacetyl exposure additionally induced expression of uncharacterised oxidoreductases (*F20G2.1*, *C06E4.3*, *F56D5.3*, *D1054.8*) and *argk-1*, an arginine kinase involved in maintaining ATP levels in invertebrates and acting as a longevity effector downstream of S6K inhibition^47^. Critically, we showed that diacetyl only boosts starvation-dependent FMO-2 (**Fig. 3e**), excluding a generalised oxidative stress effect. This was a surprising result, as chronic exposure to food cues has been reported to repress the longevity benefits of DR^48, 32^. A potential explanation is that repeated exposure to volatiles in the absence of food deteriorates the way worms perceive these as food cues. Previous studies have demonstrated that worms pre-exposed to diacetyl in the absence of food lose their attraction to diacetyl-producing lactic acid bacteria^8^. Recent results have also shown that the worm’s ability to sense and move towards volatiles, including diacetyl, declines when worms are aged on OP50 *E. coli*.^49^. This effect can be delayed by DR, which may explain why chronic odorant exposure particularly affects DR lifespan^48, 32^. We show that a single, acute exposure to diacetyl is sufficient to boost *fmo-2* expression and downstream phenotypes. We validate this finding by RNA sequencing, with a transcriptional reporter, qPCR, as well as metabolic and stress response signatures (**Fig. 3**). It remains to be determined which target tissues and sensing mechanisms are responsible for the diacetyl-induced effects reported here, a question that necessitates additional investigation. While diacetyl induces *fmo-2* over a concentration range well-established as chemoattractive (0.1% to 1%) (**Fig. 3f**) and canonically sensed by ODR-10, we show that neither the ODR-10 nor SRI-14 canonical receptors are required for *fmo-2* induction by diacetyl (**Fig. S3d**). This agrees with previous findings that the lifespan effects of chronic diacetyl exposure are independent of these receptors^24^. Blocking multiple pathways related to food sensing, neuronal signal transmission and cilia development also failed to stop the diacetyl response (**Fig. S3e - g**), pointing to a non-canonical sensing mechanism.

While it seems counterintuitive that a food-related signal would trigger a starvation marker, our findings indicate that diacetyl first induces the DHAP-Glycerol shunt, a response associated with feeding. We propose that *fmo-2* expression is a secondary effect resulting from shunt activation, when this is uncoupled from food intake. Supporting this, we show that *fmo-2* expression is also boosted when the shunt is triggered by hyperosmotic stress (**Fig. 4f - g**). While we do not identify the precise molecular mechanism linking shunt activation to FMO-2, we speculate that the diversion of glycolytic intermediates into the shunt likely plays a role in this process, potentially by depleting cellular energy levels (**Fig. 4k**) and inducing HLH-30 dependent gene expression (**Fig. 4l**). These findings are consistent with previous research published on the glycerol shunt (G3PP). Overexpression of G3PP *in vitro* decreases ATP production in rat cells^18^, and overexpression of the ortholog *pgph-2* in *C. elegans* activates HLH-30 and mimics beneficial effects of DR^50^.

Ultimately, this complex cascade may also serve to alter the animal’s behavior. Recent research has revealed novel FMO-2 functions beyond stress responses. FMO-2 regulates sensory perception, satiety, and exploratory behaviors^34^, including chemotaxis towards diacetyl and ethanol^51^, both of which we show induce its expression. The upregulation of *fmo-2* by volatile food cues may encourage the seeking of nearby bacteria, which is consistent with our finding that diacetyl exposure increases fasting-related appetite (**Fig. 3k - l**). In ecological settings, the induction of *fmo-2* by diacetyl would likely be a transient phenomenon, rapidly attenuated as the animals begin to feed.

In conclusion, we characterise a novel example of anticipatory food response directly linking a bacterial volatile to rapid gene expression and metabolic responses in food-deprived *C. elegans*. We identify the DHAP-Glycerol shunt as a crucial node where responses to feeding, starvation and hyperosmotic stress intersect.

## Methods

### *C. elegans* strains and culture

All *C. elegans* strains were maintained at 20 °C on nematode growth medium (NGM) agar plates seeded with the *Escherichia coli* (E. coli) strain OP50 unless indicated otherwise^52^.

| Strain | Genotype | Source |
| --- | --- | --- |
| Bristol N2 | Wild-type | CGC |
| VP198 | *kbls5 [gpdh-1p::GFP + rol-6(su1006)]* | CGC |
| KAE28 | *unc-119(ed3) III; seaEx13 [fmo-2p::GFP + unc-119(+)]* | CGC |
| VC1668 | *fmo-2 (ok2147) IV* | CGC |
| CX3410 | *odr-10 (ky225) X* | CGC |
| VC2123 | *sri-14 (ok2865) I* | CGC |
| CB450 | *unc-13 (e450) I* | CGC |
| CB928 | *unc-31 (e928) IV* | CGC |
| MT15434 | *tph-1 (mg280) II* | CGC |
| MT15620 | *cat-2 (n4547) II* | CGC |
| JT204 | *daf-12 (sa204) X* | CGC |
| JT6924 | *daf-19 (m86) II; daf-12 (sa204) X* | CGC |

Glucose and sorbitol plates were prepared by adding filter-sterilized glucose and sorbitol solutions to NGM agar after autoclaving, to reach a final concentration of 111 mM. Plates were seeded with live OP50 and worms were raised at 25 °C to measure *gpdh-1p::GFP* induction.

Food deprivation was performed by washing worms onto unseeded NGM plates.

#### Diacetyl exposure

Unless specified otherwise, day 1 adult worms were washed for 1 hour with M9 in rotating falcon tubes to remove bacteria, then transferred to unseeded 10 cm NGM agar plates and allowed to air dry for a few minutes. Worms were split into control (no volatile) and diacetyl-exposed plates. A 1% solution of diacetyl (2,3-Butanedione, Sigma-Aldrich) in 100% ethanol was prepared immediately before the experiment. 50 μL of this solution was pipetted on a 90 mm square of blue roll placed on the centre of the lid, then immediately closed, sealed with tape and kept lid-side down throughout the exposure.

#### RNA extraction

Worms were washed in M9 and snap-frozen in liquid nitrogen. The pellet was resuspended in TRI Reagent (Zymo) and RNA extraction was performed using the Direct-zol RNA MicroPrep Kit (Zymo Research) according to the manufactureŕs recommendations.

#### RNA sequencing

Three biological replicates were collected and RNA extracted as described above. Poly-A selected cDNA libraries were produced using the NEBNext Ultra II Directional RNA Library Kit and sequenced on the Illumina NextSeq2000 platform. Raw sequencing data (FASTQ files) were processed using the nf-core/rnaseq pipeline (version 3.12.0) executed via Nextflow (version 23.04.2)^53,54^. Raw sequencing reads were subjected to quality trimming and adapter removal using Trim Galore^55^ (version 0.6.7) with Cutadapt (version 3.4). Comprehensive quality control reports summarizing all initial and post-alignment metrics (e.g., read distribution, coverage, duplication rates) were generated using MultiQC^56^. Reads were mapped to the reference genome and transcriptome using STAR^57^ (version 2.7.9a for indexing, 2.7.10a for alignment). Transcript-level quantification was performed using Salmon (version 1.10.1) via a quasi-mapping approach^58^. Separately, gene-level counts were generated using featureCounts (Subread package version 2.0.1). Reads aligned to the genome by STAR were further visualised and quantitated using SeqMonk^59^ (v1.48.0). Limma and intensity difference filters were combined to define differentially expressed genes (P < 0.005). Tissue expression scores for genes upregulated by diacetyl were calculated using an online tool^60^ available at “https://worm.princeton.edu/”. RNA sequencing data generated during this study have been deposited in the NCBI Gene Expression Omnibus (GEO) under accession number GSE310881.

#### Metabolomics and lipidomics

Worms were washed in M9 and exposed to diacetyl on unseeded plates as described above. Worms were exposed to diacetyl for one or three hours, then snap-frozen. For the fed condition, worms were directly snap-frozen before M9 starvation. For hyperosmotic stress, worms were transferred to 200 mM NaCl seeded plates for three hours before snap-freezing. Experiments were carried out with ten biological replicates.

Frozen worm pellets were extracted overnight using ice-cold LC–MS grade solvent consisting of 800 µL 80% methanol / 20% water and 700 µL 53:33:12:1 (MeOH:CHCl₃:H₂O:acetic acid, v/v/v/v). Isotopically labeled internal standards prepared in 80% methanol / 20% water were spiked into each sample proportional to the average number of worms per condition. Samples were sonicated in a water bath to dissociate worms, vortexed, and incubated overnight at −80 °C. The extracts were then centrifuged at 30,000 × g for 20 min at 4 °C, and the supernatant was collected and dried in a speed vacuum concentrator at 4 °C overnight. Dried samples were resuspended in ACN:MeOH:H₂O (40:40:20, v/v/v) at volumes normalised to worm number, followed by centrifugation at 30,000 × g for 20 min at 4 °C. The clarified supernatants were transferred to glass vials arranged in a 96-well autosampler plate. Throughout processing, samples were handled on dry ice, and during LC–MS analysis they were maintained at 4 °C in the autosampler.

Metabolites were measured using an Exploris 240 mass spectrometer (Thermo Fisher Scientific) connected to a Vanquish Duo UHPLC system (Thermo Fisher Scientific) operated in both positive and negative ion modes. Anionic metabolites (negative ion mode) were separated on an Acquity Premier GLY BEH 130 Å, 1.7 µm, 2.1 × 150 mm column (Waters, #186009524) at a flow rate of 0.40 mL/min using a binary gradient consisting of Mobile Phase A (10 mM ammonium acetate in 90:10 H₂O:acetonitrile with 5 µM medronic acid) and Mobile Phase B (10 mM ammonium acetate in 80:10 acetonitrile:H₂O with 5 µM medronic acid). The gradient started at 0% Mobile Phase A and was increased to 30% A over 8 minutes, followed by 1 minute at 50% A. At 9.5 minutes, the gradient was returned to 0% A for 50 seconds and then held at 0% A for an additional 3 minutes. The column temperature was maintained at 40 °C. Cationic metabolites (positive ion mode) were separated on an InfinityLab Poroshell 120 HILIC-Z column, 2.1 × 150 mm, 2.7 µm (Agilent Technologies, #683775-924) at a flow rate of 0.40 mL/min with Mobile Phase A (10 mM ammonium formate in water with 0.1% formic acid) and Mobile Phase B (acetonitrile with 0.1% formic acid). The gradient started at 5% A and was held for 1 minute, then increased linearly to 50% A over 6 minutes. The mobile phase was held at 50% A for 3 minutes before being returned to 5% A and held for 2 minutes. The column temperature was maintained at 40 °C. Full-scan MS data were acquired under the following conditions: m/z range 70–900; resolving power 120,000 (at m/z 200); automatic gain control target set to Standard; and maximum ion injection time set to Auto. Ion source parameters were: spray voltage, static; ion transfer tube temperature, 275 °C; RF lens level, 75; heater temperature, 320 °C; sheath gas, 50 (arbitrary units); and auxiliary gas, 10 (arbitrary units). Raw LC–MS data were imported into Skyline, and chromatographic peaks were manually inspected and curated for peak picking and integration to generate peak area values for downstream quantitative analysis.

Lipids were measured using a SCIEX Triple Quad 7500 mass spectrometer (SCIEX) operated in LC–MS/MS mode and connected to a Vanquish Duo UHPLC system (Thermo Fisher Scientific) operated in both positive and negative ion modes. Lipids in negative ion mode were separated on an ACQUITY UPLC BEH C18 column, 130 Å, 1.7 µm, 2.1 × 100 mm (Waters Technologies Corporation, #186002352) at a flow rate of 0.25 mL/min using a binary gradient consisting of Mobile Phase A (10 mM ammonium acetate in 60:40 acetonitrile:water) and Mobile Phase B (10 mM ammonium acetate in 90:10 isopropanol:acetonitrile). The gradient started at 45% Mobile Phase B and was increased to 99% B over 8 minutes and held at 99% B for 1 minute. At 9 minutes, the gradient was returned to 55% B for 1 minute and then held at 55% B for an additional 2 minutes. The column temperature was maintained at 55 °C. Lipids in positive ion mode were separated on the same ACQUITY UPLC BEH C18 column (130 Å, 1.7 µm, 2.1 × 100 mm; Waters Technologies Corporation, #186002352) at a flow rate of 0.25 mL/min using a binary gradient consisting of Mobile Phase A (10 mM ammonium formate in 60:40 acetonitrile:water with 0.1% formic acid) and Mobile Phase B (10 mM ammonium formate in 90:10 isopropanol:acetonitrile with 0.1% formic acid). The gradient started at 45% Mobile Phase B and was increased to 99% B over 8 minutes and held at 99% B for 1 minute. At 9 minutes, the gradient was returned to 55% B for 1 minute and then held at 55% B for an additional 2 minutes. The column temperature was maintained at 55 °C. Data acquisition was performed in scheduled multiple reaction monitoring (MRM) mode using an optimised MS/MS transition and retention time table. Source and gas parameters were as follows: ion source gas 1, 45 psi; ion source gas 2, 60 psi; curtain gas, 50 psi; collision gas (CAD), 9 (arbitrary units); source temperature, 500 °C; and ion spray voltage, 4200 V. Raw LC–MS/MS data were imported into Skyline, and chromatographic peaks were manually inspected and curated for peak picking and integration to generate peak area values for downstream quantitative analysis.

Chromatographic peak integration and extraction from raw data were performed in a targeted manner using Skyline^61^. Metabolites with poor peak shapes were excluded. Analyte measures were normalised using probabilistic quotient normalisation^62^. Briefly, analytes (excluding spiked internal standards) were median-scaled across samples. The median of these median-scaled analyte values across all analytes was used for each sample as the normalisation factor. Each analyte measurement value was divided by this normalisation factor to compute the final normalised value. Quality-control replicates from pooled samples were interspersed with study samples. Analytes with coefficient of variation (CV) values across these replicates that exceeded 25% after normalisation were removed. In cases where an analyte was measured across multiple LC-MS methods, measurements from the LC-MS method that produced the lower CV value for that analyte were used in downstream analyses. Differential abundance testing was performed on normalised and log transformed measurements using Welch’s T-Test. Adjustment for multiple comparisons was completed using the Benjamini & Hochberg approach. Analyte ratios for ATP/ADP were computed using the unnormalised LC-MS peak areas.

#### Glycerol measurements

Worms were exposed to diacetyl for two hours as described above and snap-frozen. Eight biological replicates were collected, frozen worms pellets were resuspended in RIPA buffer (150 mM NaCl, 1% NP40, 0.5% sodium deoxycholate, 0.1% sodium dodecyl sulfate (SDS), 50 mM Tris-HCl, pH 8.0, completed with protease inhibitors), sonicated and spun down. Protein quantification was done by bicinchoninic acid assay (Pierce BCA Protein Assay Kit, Thermo Fisher). Glycerol concentration was measured with a glycerol assay kit (MAK117, Sigma-Aldrich) following manufactureŕs protocol. Samples were pre-diluted to fall in the detection range of the colorimetric detection. Glycerol concentration was normalized by the protein concentration in each sample. Error bars represent means ± SD and were analysed by Welch’s t-test.

#### Acute hyperosmotic stress resistance assay

Worms were exposed to diacetyl for one hour as described above. Worms were washed off the exposure plates in M9 and spinned down. 30 μL of worm pellets were pipetted on unseeded high salt plates containing 500 mM NaCl. Control and diacetyl exposed worms were placed in droplets side to side to record under the same field of view. Starting three minutes after pipetting, one minute videos were recorded on a Leica S9i stereo microscope. For quantification of hyperosmotic stress resistance, body bends of each worm were manually recorded from the last 30 seconds of each video. Three biological replicates were performed per experiment, with at least 15 worms scored per condition in each replicate. Error bars represent means ± SD and were analysed by Mann-Whitney test or one-way ANOVA followed by Sidak’s post hoc test as indicated.

#### Worm imaging

Worms were arranged in stacks on unseeded NGM plates and kept on ice. Images were taken on a Leica M205 FCA fluorescence stereo microscope. Images were acquired with the Leica Application Suite X. Images were quantified with ImageJ (version 2.9.0). Scale bar is indicated in the figure legends. Three biological replicates were performed per experiment, with at least 15 worms imaged/quantified per condition in each replicate. For imaging of *fmo-2p::GFP* reporter activation by diacetyl, worms were washed off unseeded exposure plates and transferred to seeded NGM plates for 2 hours before imaging to stop reporter induction and allow GFP maturation.

#### qPCR

Gene expression levels were quantified using a 1-step RT-qPCR kit (A6020, Promega) on a QuantStudio 7 Pro System (Applied Biosystems). Expression levels of the genes *act-1* and *tba-1* were used as internal controls for normalization. At least three biological replicates were analysed per experiment.

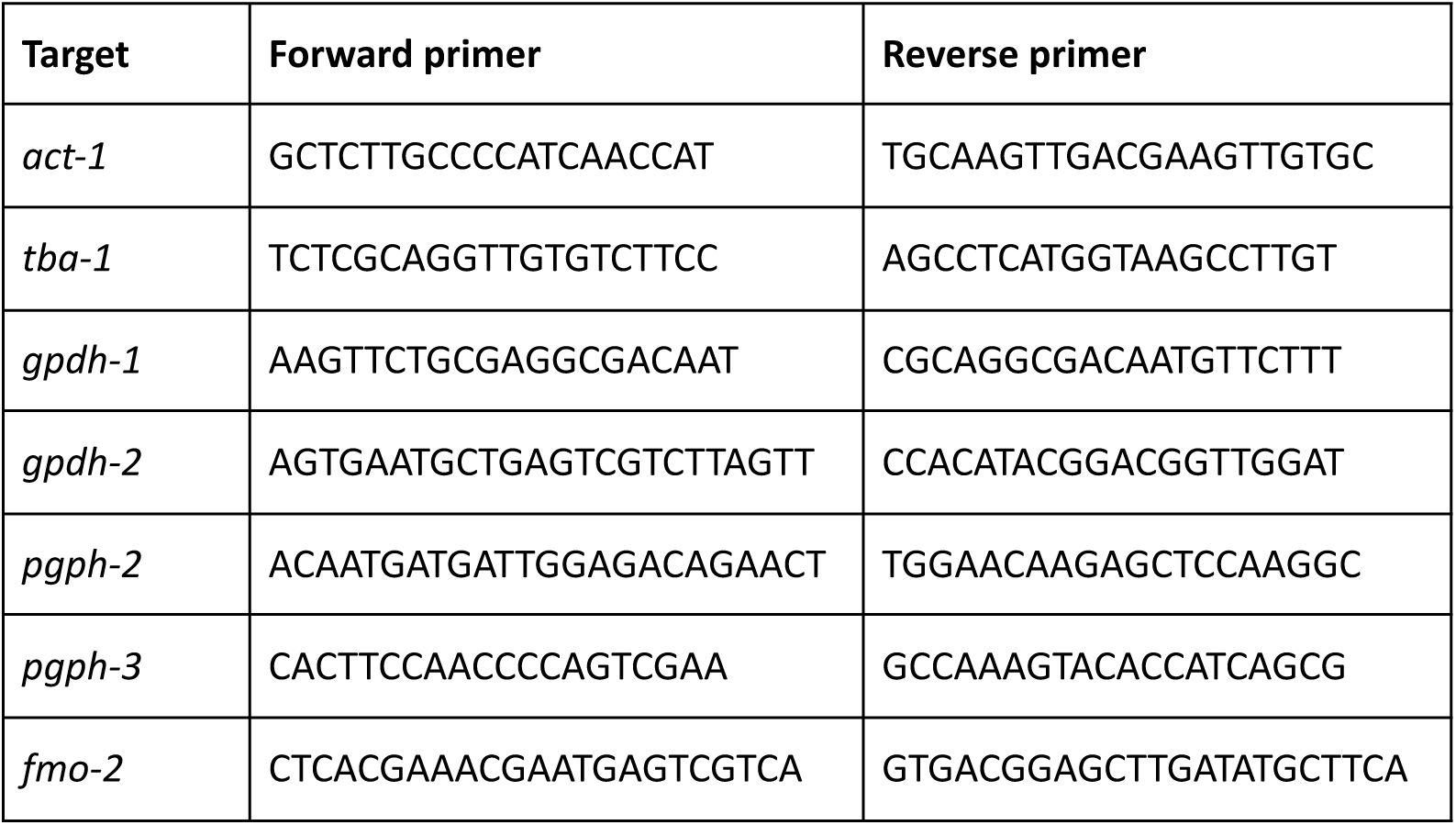

#### Ivermectin treatment

For ivermectin treatment of the *fmo-2p::GFP* reporter strain, worms were treated as described above but 100 nM of Ivermectin (I8898, Sigma-Aldrich) was added to the M9 during the one hour wash preceding diacetyl exposure.

#### Feeding rate measurements

Worms were exposed to diacetyl for three hours as described above, then immediately washed onto plates seeded with GFP-expressing OP50 (CGC) and allowed to feed for five minutes. Worms were then washed onto unseeded NGM plates kept on ice for imaging. Three biological replicates were performed per experiment, with 15 worms quantified per condition in each replicate. Error bars represent means ± SD and were analysed by Brown-Forsythe and Welch ANOVA test followed by Dunnett’s T3 post hoc test.

#### Thermotolerance assays

Day 1 adult worms were starved by transferring to unseeded NGM plates for three hours. For combination with diacetyl exposure worms were treated as described above. For combination with hyperosmotic stress, worms were pre-treated for three hours on seeded NGM plates with 200 mM NaCl. Worms were maintained on seeded plates after day 1 treatments. On day 2, worms were placed at 36 °C for three hours and then recovered at 20 °C overnight. Survival was scored on day 3 for (touch-provoked) movement. Throughout the experiment, strain and/or treatment were unknown to the researcher. Three biological replicates were performed per experiment, with at least 100 worms scored per condition in each replicate. Error bars represent means ± SD and were analysed by ordinary one-way ANOVA followed by Sidak’s post hoc test.

#### RNAi experiments

For RNAi-mediated knockdown of specific genes, HT115 bacteria carrying vectors for dsRNA of the target gene under a promoter inducible by isopropyl β-d-1-thiogalactopyranoside (IPTG) and ampicillin resistance were used. Bacteria were seeded on NGM plates containing 100 µg/µL ampicillin (Merck Millipore) and 1 mM IPTG (Roth). Eggs were transferred to RNAi plates. RNAi against luciferase was used as nontargeting control. All RNAi clones were obtained from the Ahringer and Vidal RNAi libraries^63, 64^. Clones were validated by plasmid purification (Zyppy Plasmid Miniprep Kit, Zymo research) and whole plasmid sequencing was performed by Plasmidsaurus using Oxford Nanopore Technology with custom analysis and annotation.

#### Hyperosmotic stress development

Eggs were placed on either control RNAi plates or RNAi plates supplemented with 200 mM NaCl. Development was assessed after three days and worm area was quantified with ImageJ (version 2.9.0). Three biological replicates were performed per experiment, with 15 worms measured per condition in each replicate. Error bars represent means ± SD and were analysed by ordinary one-way ANOVA followed by Dunnett’s post hoc test.

#### Hyperosmotic stress survival

Worms were grown on RNAi plates as described above and transferred on day 1 of adulthood to RNAi plates supplemented with 400 mM NaCl. Survival was assessed two and four days after the start of hyperosmotic stress by monitoring (touch-provoked) movement and pharyngeal pumping. Worms that had undergone internal hatching, vulval bursting, or worms crawling off the plates were censored. Throughout the experiment, strain and/or treatment were unknown to the researcher. Three biological replicates were performed per experiment, with 100 to 300 worms scored per condition in each replicate.

## Supplementary figures

**Supplementary Figure 1.**
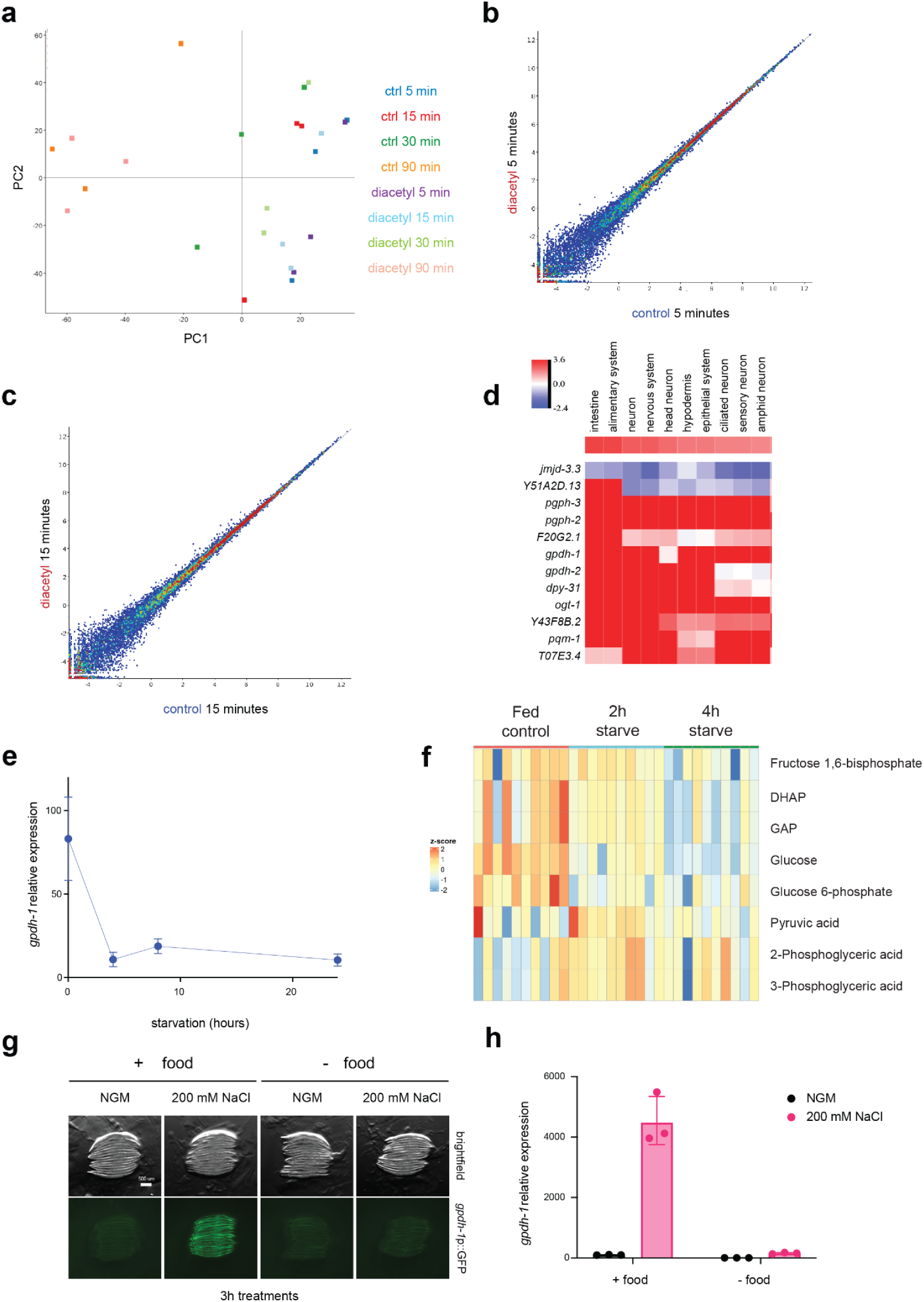
**a**, PCA plot of diacetyl transcriptomic time course from Figure 1a. **b** - **c,** Scatter plots comparing gene expression between control and diacetyl groups at 5 (**b**) and 15 (**c**) minutes of exposure. **d**, Predicted tissue expression scores for genes upregulated by diacetyl, in the top ten enriched tissues. **e,** *gpdh-1* expression levels over twenty-four hours of food deprivation. Measured by qPCR, normalised to 0h time point. Error bars represent means ± SD, n = 3. **f**, Heatmap comparing levels of glycolytic intermediates detected from metabolomics in fed condition and after two or four hours of food deprivation (n = 10). **g**, Representative images of *gpdh-1* transcriptional reporter worms exposed to hyperosmotic stress (200 mM NaCl), in the presence or absence of food. Images shown are from one experiment out of three biological replicates. Additional replicates are available in Figure 1 - Source data S1g. **h**, *gpdh-1* expression levels at control osmolarity (NGM) or 200 mM NaCl (3 hours), in the presence or absence of food (n = 3). Measured by qPCR, normalised to NGM fed levels.

**Supplementary Figure 2.**
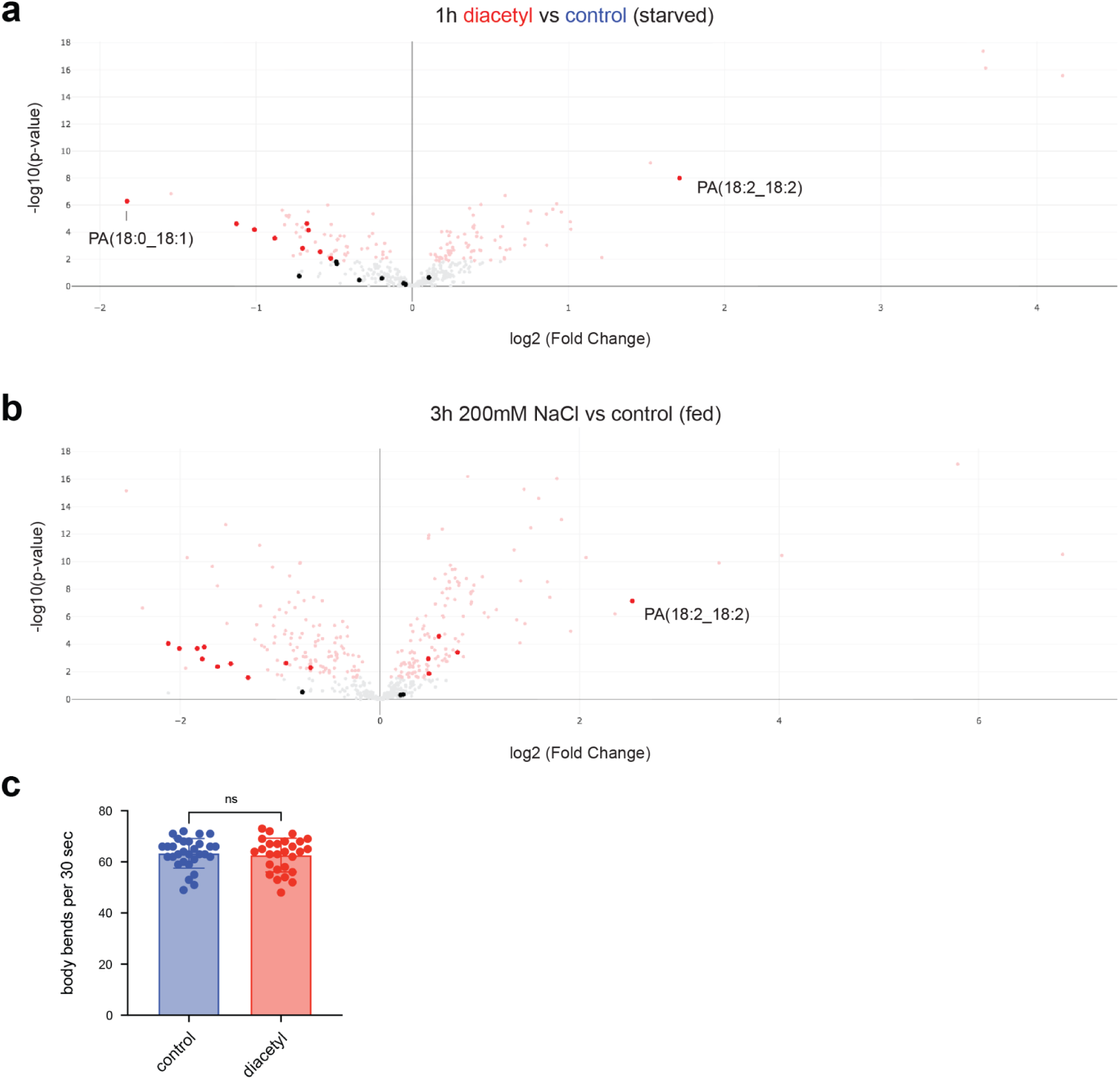
**a**, Volcano plot comparing metabolomic effects of diacetyl exposure (one hour) to control food deprivation. Metabolites significantly altered in red (FDR <0.05, Welch’s t-test). Highlighted are metabolites classified as phosphatidic acids (PAs). **b,** Volcano plot comparing metabolomic effects of hyperosmotic stress (three hours, 200 mM NaCl plates with food) to control NGM plates. Metabolites significantly altered in red (FDR <0.05, Welch’s t-test). Highlighted are metabolites classified as PAs. **c,** Effects of diacetyl pre-exposure on thrashing rates. Worms were pre-exposed to diacetyl for one hour (in the absence of food), before thrashing was recorded on unseeded NGM plates. Data is pooled from two biological replicates; n = 30 worms in the control group; n = 27 in the diacetyl group. Error bars represent means ± SD, unpaired t-test with ns: not significant.

**Supplementary Figure 3.**
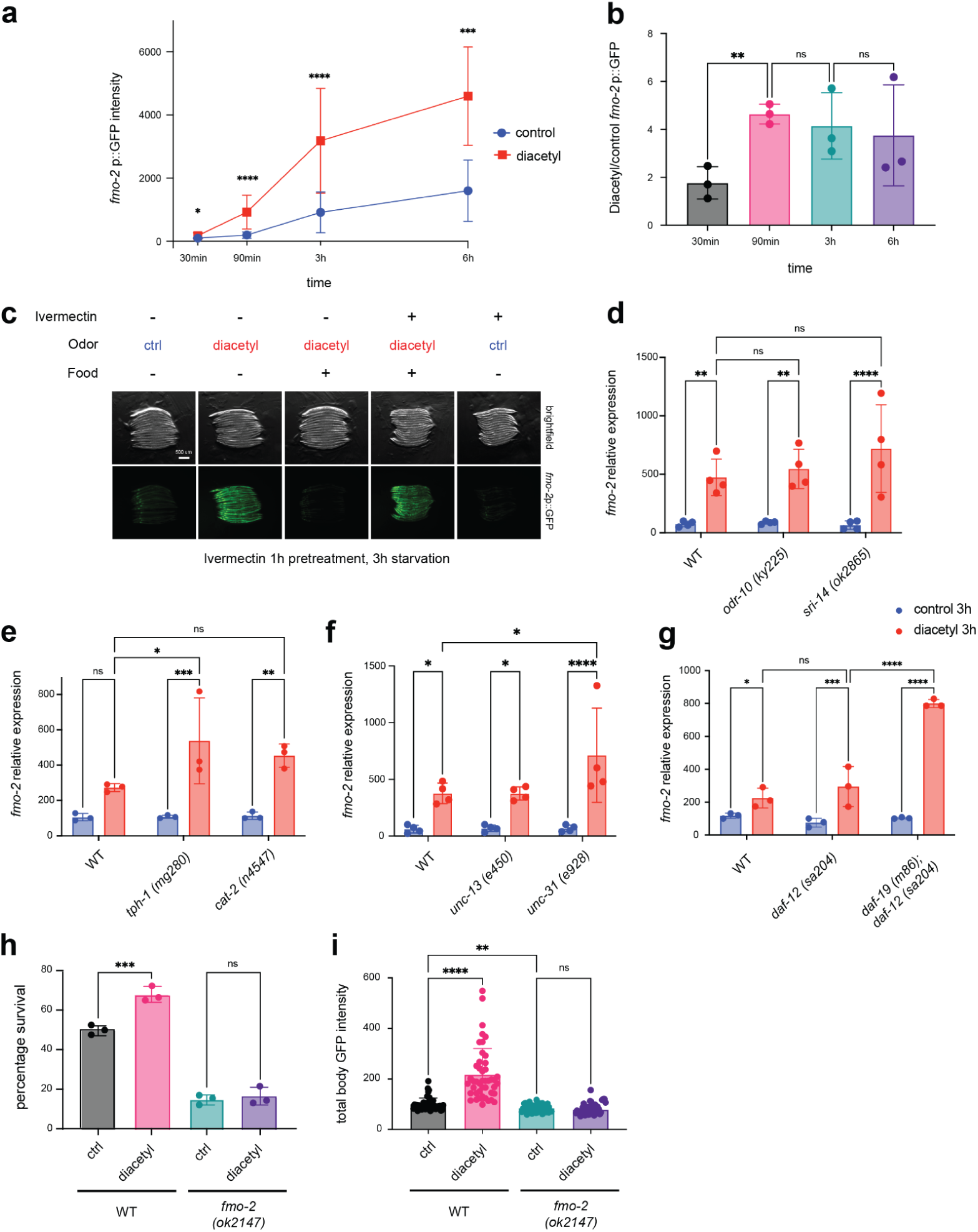
**a**, Quantification of *fmo-2* reporter induction in experiment from Figure 3d. Data is pooled from all three replicates in Figure 3 - Source data d (n = 45 worms per condition). Error bars represent means ± SD, Kruskal-Wallis test followed by Dunn’s post hoc test with *P = 0.0347 (30 min), ****P < 0.0001, ***P = 0.0002 (6h). **b**, Quantification from (a) with diacetyl *fmo-2* reporter induction normalised to the control starvation for each timepoint. Error bars represent means ± SD, RM-one way ANOVA followed by Tukey’s post hoc test with **P = 0.0074, ns: not significant. **c**, Representative images of *fmo-2* reporter worms treated with ivermectin for one hour, followed by diacetyl exposure in the presence or absence of food for three hours. Images shown are from one experiment out of three biological replicates. Additional replicates are available in Figure 3 - Source data e. **d** - **g**, *fmo-2* expression levels in worms starved either in control conditions or in the presence of diacetyl for three hours. Detected by qPCR. Normalised to control for each genotype. **d,** *fmo-2* expression levels in WT, odr-10 (ky225) and sri-14 (ok2865). Error bars represent means ± SD (n = 4), two-way ANOVA followed by Tukey’s post hoc test with **P = 0.0059 ctrl vs diacetyl (WT), **P = 0.0021 ctrl vs diacetyl (odr-10), ****P < 0.0001 ctrl vs diacetyl (sri-14). **e,** *fmo-2* expression levels in WT, tph-1 (mg280) and cat-2 (n4547). Error bars represent means ± SD (n = 3), two-way ANOVA followed by Tukey’s post hoc test with ***P = 0.0003 ctrl vs diacetyl (tph-1), **P = 0.0017 ctrl vs diacetyl (cat-2), *P = 0.0223 WT vs tph-1 (diacetyl). **f,** *fmo-2* expression levels in WT, unc-13 (e450) and unc-31 (e928). Error bars represent means ± SD (n = 4), two-way ANOVA Tukey’s post hoc test with *P = 0.0211 ctrl vs diacetyl (WT), *P = 0.0244 ctrl vs diacetyl (unc-13), ****P < 0.0001 ctrl vs diacetyl (unc-31), *P = 0.0375 WT vs unc-31 (diacetyl). **g,** *fmo-2* expression levels in WT, daf-12 (sa204) and daf-12; daf-19 (sa204; m86). Error bars represent means ± SD (n = 3), two-way ANOVA Tukey’s post hoc test with *P = 0.041 ctrl vs diacetyl (WT), ***P = 0.0005 ctrl vs diacetyl (daf-12), ****P < 0.0001 ctrl vs diacetyl (daf-12; daf-19), ****P < 0.0001 daf-12 vs daf-12; daf-19 (diacetyl). **h**, Survival (day 3 of adulthood) of worms exposed for three hours to diacetyl in the absence of food (day 1) and heat-shocked on day 2. Error bars represent means ± SD (n = 3), ordinary one-way ANOVA followed by Sidak’s post hoc test with ***P = 0.004. **i**, Quantification of GFP intensity in worms fed with GFP expressing bacteria following a three-hour exposure to diacetyl in the absence of food. Data is pooled from three replicates in Figure 3 - Source data S3i (n = 45 worms per condition). Error bars represent means ± SD, Brown-Forsythe and Welch ANOVA test followed by Dunnett’s T3 post hoc test, ****P < 0.0001, **P = 0.0011.

**Supplementary Figure 4.**
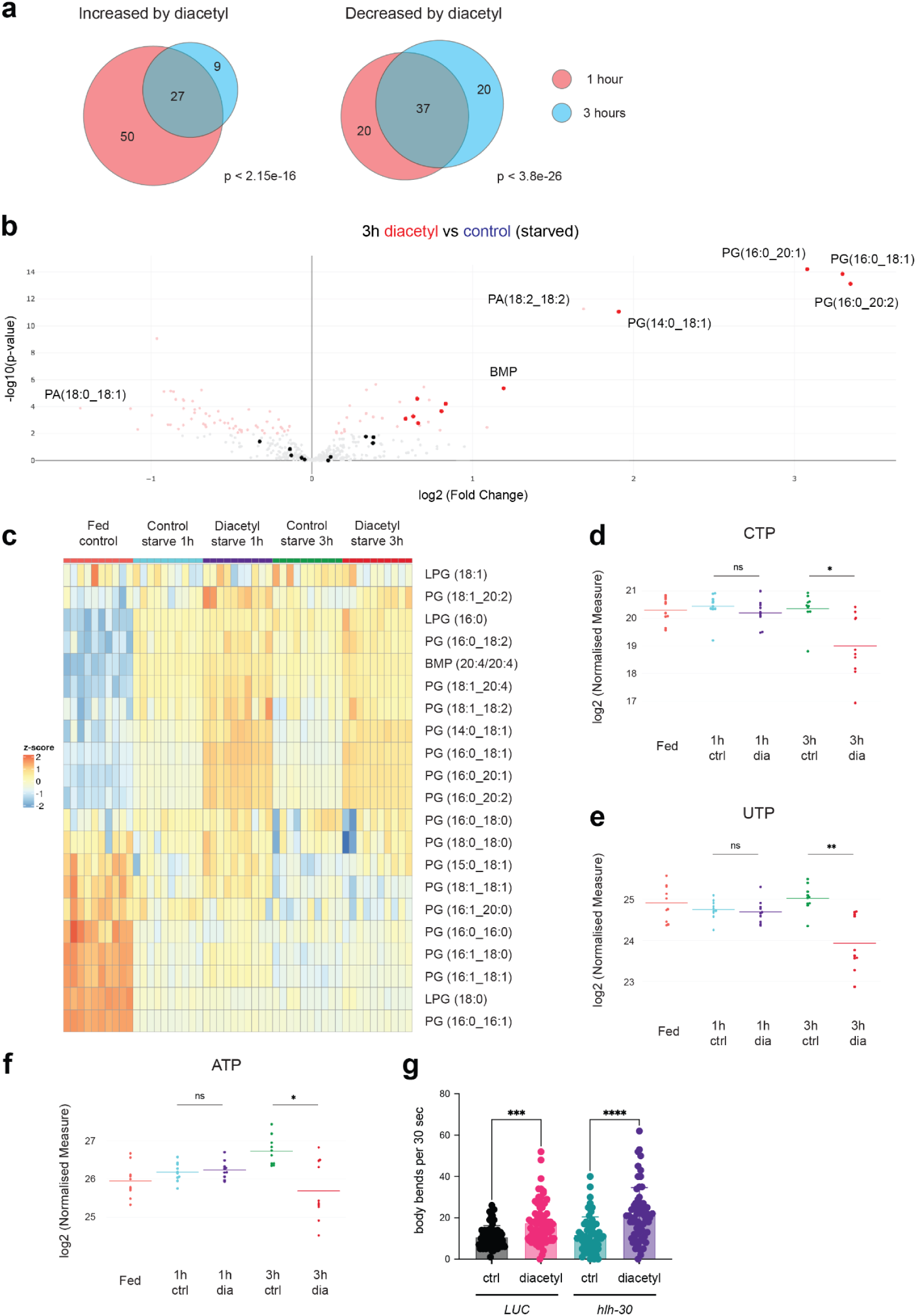
**a**, Overlap between metabolites significantly altered (FDR <0.05, Welch’s t-test) by diacetyl, after one or three hours exposure in the absence of food. Significance determined by hypergeometric test; increased metabolites x = 27, N = 472, p < 2.15e-16; decreased metabolites x = 37, N = 472, p < 3.8e-26. **b**, Volcano plot comparing metabolomic effects of diacetyl exposure (three hours) to control food deprivation. Metabolites significantly altered in red (FDR <0.05, Welch’s t-test). Highlighted are metabolites classified as phosphatidylglycerols (PGs). **c,** Heatmap comparing levels of all phosphatidylglycerols (PGs) in fed condition, control food deprivation (control, one and three hours) and diacetyl exposure during food deprivation (diacetyl, one and three hours) (n = 10). **d** - **f**, Individual metabolite levels in fed and food deprived worms, starved either in control (ctrl) conditions or with diacetyl (dia) exposure. One hour and three hour time points. Log2 normalised levels on vertical axes (n = 10). **d,** CTP levels, Welch’s t-test with *P = 0.03. **e,** UTP levels, Welch’s t-test with **P = 0.006. **f,** ATP levels, Welch’s t-test with *P = 0.02. **g,** Effects of diacetyl pre-exposure on acute hyperosmotic stress resistance. Worms raised on non-targeting Luciferase (LUC) RNAi targeting *hlh-30* were pre-exposed to diacetyl for one hour (in the absence of food), before thrashing was recorded on hyperosmotic stress plates (500 mM NaCl). Data is pooled from three biological replicates; n = 64 worms (LUC, ctrl), n = 72 (LUC, diacetyl), n = 61 (*hlh-30*, ctrl), n = 64 (*hlh-30*, diacetyl). Error bars represent means ± SD, ordinary one-way ANOVA followed by Sidak’s post hoc test with ***P = 0.0001 (LUC), ****P < 0.0001 (*nhr-49*).

**Supplementary Figure 5.**
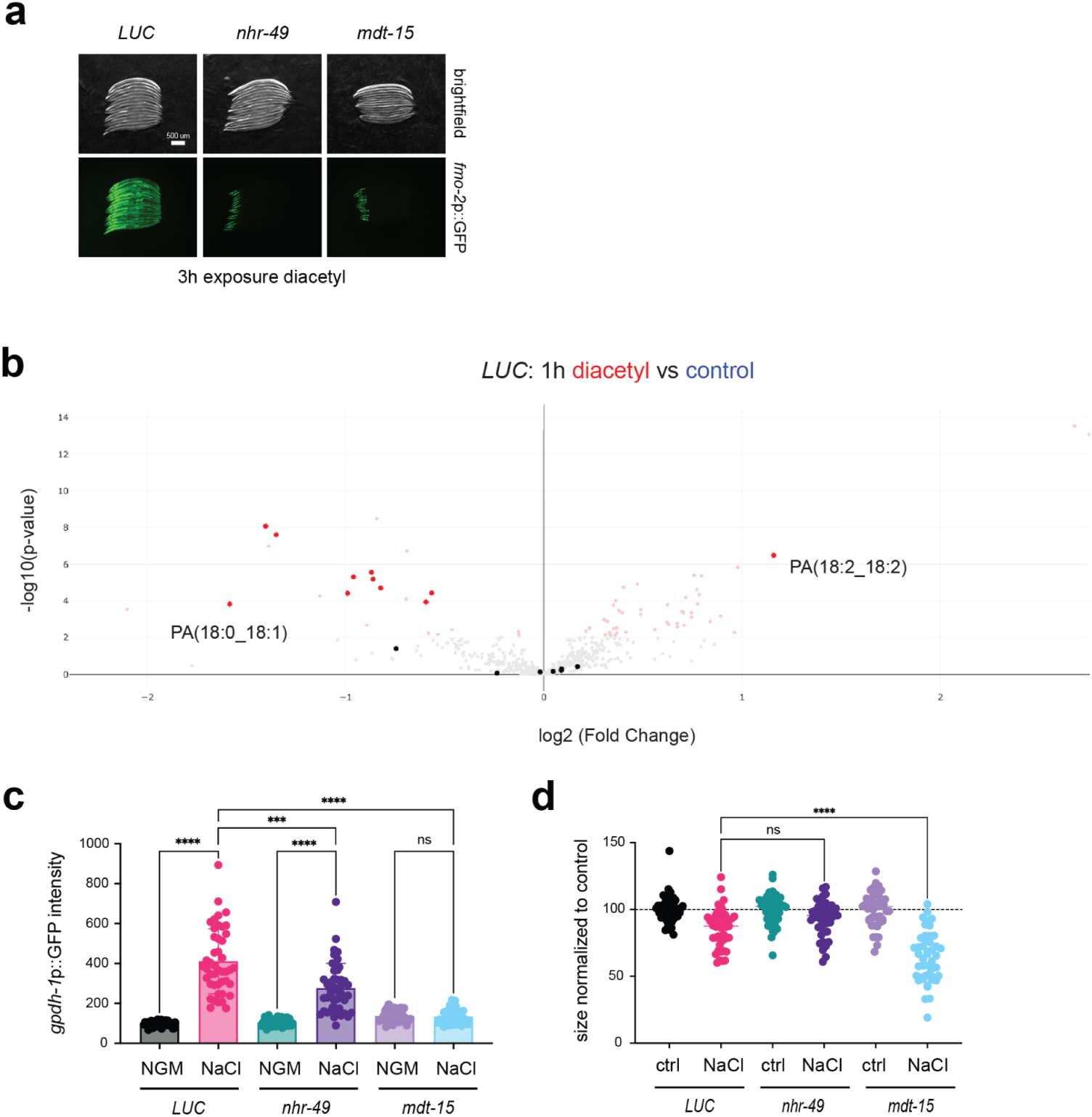
**a**, Representative images of *fmo-2* transcriptional reporter induction by diacetyl exposure in the absence of food (three hours), in worms raised on non-targeting Luciferase (LUC) or RNAi targeting *nhr-49* or *mdt-15*. Images shown are from one experiment out of three biological replicates. Additional replicates are available in Figure 5 - Source data S5a. **b**, Volcano plot comparing metabolomic effects of diacetyl exposure (one hour) to control food deprivation, in worms raised on non-targeting Luciferase (LUC) RNAi. Metabolites significantly altered in red (FDR <0.05, Welch’s t-test). Highlighted are metabolites classified as phosphatidic acids (PAs). **c**, Quantification of *gpdh-1* reporter induction in experiment from Figure 5e. Data is pooled from all three replicates (n = 45 worms per condition). Error bars represent means ± SD, Brown-Forsythe and Welch ANOVA tests followed by Dunnett’s T3 post hoc test with ****P < 0.0001 NGM vs NaCl (LUC), ****P < 0.0001 NGM vs NaCl (*nhr*-49), ***P = 0.0002 LUC vs *nhr*-49 (NaCl), ****P < 0.0001 LUC vs *mdt-15* (NaCl), ns: not significant. **d**, Quantification worm size in experiment from Figure 5f. Normalised to size of worms in control osmolarity for each RNAi condition. Ordinary one-way ANOVA followed by Dunnett’s post hoc test with ****P < 0.0001.

